# Activity regulation of a glutamine amidotransferase bienzyme complex by substrate-induced subunit interface expansion

**DOI:** 10.1101/2025.01.26.634917

**Authors:** Franziska Jasmin Funke, Sandra Schlee, Isabel Bento, Gleb Bourenkov, Reinhard Sterner, Matthias Wilmanns

**Affiliations:** Institute of Biophysics and Physical Biochemistry, Regensburg Center for Biochemistry, University of Regensburg, 93040 Regensburg, Germany; European Molecular Biology Laboratory, Hamburg Unit, 22607 Hamburg, Germany; University Hamburg Clinical Center Hamburg-Eppendorf, 20251 Hamburg, Germany

**Author notes:** **Corresponding Authors** Reinhard Sterner, Matthias Wilmanns.

**Keywords:** catalysis synchronization, multienzyme complex, nitrogen containing metabolites, ammonia utilization in catalysis, substrate/product sequestration, enzyme product/substrate tunnel, active site interface expansion

## Abstract

Glutamine amidotransferases are multienzyme machineries in which reactive ammonia is generated by a glutaminase and then transferred through a sequestered protein tunnel to a synthase active site for incorporation into diverse metabolites. To avoid wasteful metabolite consumption, there is a requirement for synchronized catalysis but any generally applicable mechanistic insight is still lacking. As synthase activity depends on glutamine turnover, we investigated possible mechanisms controlling glutaminase catalysis, using aminodeoxychorismate synthase involved in folate biosynthesis as a model. By analyzing this system in distinct states of catalysis, we found that incubation with glutamine leads to a subunit interface expansion by one third of its original area. These changes completely enclose the glutaminase active site for sequestered catalysis and subsequent transport of volatile ammonia to the synthase active site. In view of similar rearrangements in other glutamine amidotransferases, our observations provide a general mechanism for catalysis synchronization of this multienzyme family.

## INTRODUCTION

One of the dogmas in enzymology is in the requirement for conformational plasticity of the active site during catalysis to allow substrate access, turnover and product release ^1^. In multienzyme systems, in which the product of the first catalyzed reaction is subsequently transferred through a protein tunnel to become the substrate of the second enzyme, there are additional needs for sequestration to prevent its leakage during transport. These systems are subject to multiple mechanisms of activity regulation, due to the additional requirement to synchronize their activities to avoid wasteful consumption of metabolites ^2–4^. An architecturally and mechanistically rich family of such multienzyme systems is represented by glutamine amidotransferases (GATs) that are responsible for the incorporation of nitrogen within numerous metabolic pathways ^5–7^. GATs comprise minimally a glutaminase that produces ammonia, which reacts with a synthase-bound substrate to yield specific nitrogen-containing products. Since ammonia is mostly converted into the inert protonated ammonia ion under physiological pH conditions, in these GATs there is a strict requirement for sequestered transfer of reactive ammonia from the glutaminase active site to the synthase active site. Because of the volatility of ammonia even small defects in sequestration can lead to substantial leakage and uncoupling of GAT activities ^8–11^.

Whereas GAT synthases are structurally and functionally diverse, known GAT glutaminases are restricted either to the class I α/β-hydrolase fold comprising a cysteine/histidine/glutamate catalytic triad or to the class II Ntn-hydrolase fold with an N-terminal catalytic cysteine ^6^. While the GAT synthase active sites are generally positioned in distinct and separate locations of the respective enzymatic subunits (or domains in case of fusion enzymes), the glutaminase active sites are often situated next to the interface between the glutaminase subunit and the interacting synthase subunit (or domain), adjacent to the entrance to distinct ammonia tunnels towards the synthase active site ^7,9^. Generally, GAT glutaminases are noticeably active only in presence of the synthase and are further stimulated by the synthase substrate ^8,12,13^. Since the supply of reactive ammonia is crucial for synthase activity, glutaminase activity regulation is determinant for the overall GAT catalysis.

Here, we chose aminodeoxychorismate synthase (ADCS) from *Escherichia coli*, which consists of a class I glutaminase subunit (PabA) and a synthase subunit (PabB), to investigate the mechanism of glutaminase catalysis regulation and its impact on overall GAT activity ^14^. PabB catalyzes the amination of chorismate (Cho) to 4-amino-4-deoxy-chorismate, which is a committed step in folic acid biosynthesis. It belongs to a structurally conserved group of Cho utilizing enzymes, known as menaquinone, siderophore and tryptophan (MST) enzymes, which use either water or ammonia as a nucleophile ^15,16^. These enzymes are of great interest as they are found in various pathogens and hence can be exploited as antimicrobial drug targets ^15,17^. Common to MST synthases is that their active site is deeply embedded within the respective subunit or domain with a strict requirement of substrate (Cho) co-binding in the presence of a highly conserved Mg^2+^ binding site ^15,16^. Despite knowledge of several structures and available biochemical data, only little is known about how glutaminase catalysis is regulated in these MST enzyme machineries.

We chose ADCS for mechanistic investigation due to its reported transient assembly in the absence of Gln, which was enhanced upon Gln addition ^18^. These data were suggestive of a role for Gln binding in forming a constitutive, catalytically competent ADCS complex ^19^. By a combined structural and biochemical approach, we demonstrate that Gln binding to the ADCS complex induces an expansion from a generic, substrate-independent to a substrate-induced extended interface between the two subunits PabA and PabB. This expansion leads to the completion and sequestration of a catalytically competent glutaminase active site from external solvent, which is only open to the ammonia tunnel towards the synthase active site. Comparison of our findings with data from other GATs reveals a generally applicable mechanistic principle for tunnel-mediated multienzyme complexes with highly regulated and synchronized activities.

## RESULTS

### Biochemical and structural basis of ADCS glutaminase activity regulation

To investigate the molecular mechanism of ADCS activity regulation, we characterized the enzyme complex from *Escherichia coli* structurally, biophysically, and biochemically. As evidenced by analytical gel filtration and isothermal titration calorimety (ITC) data, the PabA and PabB subunits form a heterodimeric assembly with a dissociation constant K_D_ = 0.16 ± 0.06 μM. While we found only residual glutaminase activity in separate PabA, the catalytic glutaminase efficiency k_cat_/K_M_ ^Gln^ of the ADCS complex in the presence of Mg^2+^ and Cho was 1.09 ± 0.13 mM^-1^s^-**1**^ **(Table 1)**, which is similar to previously reported data ^19,20^.

**Table 1:** Effects of PabA or PabB interface mutations on PabA-PabB assembly and glutaminase activity.

|  | Analytical SEC <sup>a</sup> |  | ITC <sup>b</sup> | Catalytic activity <sup>c</sup> |  |  |
| --- | --- | --- | --- | --- | --- | --- |
|  | -Gln | +Gln | K<sub>D</sub> [μM] | K<sub>M</sub><sup>Gln</sup> [mM] | k<sub>cat</sub> [s<sup>-1</sup>] | k<sub>cat</sub>/K<sub>M</sub><sup>Gln</sup> [mM<sup>-1</sup>s<sup>-1</sup>] |
| PabA/PabB |  |  |  |  |  |  |
| wt/wt |  |  |  | 0.34 ± 0.03 | 0.37 ± 0.010 | 1.09 ± 0.13 |
| wt/wt, -Cho | + | + | 0.16 ± 0.06 | 0.09 ± 0.01 | 0.22 ± 0.007 | 2.41 ± 0.47 |
| wt/- |  |  |  | 12.2 ± 0.54 | 4.4 *10<sup>-3</sup> ± 6.1 *10<sup>-5</sup> | 3.6 *10<sup>-4</sup> ± 2.1 *10<sup>-5</sup> |
| **A, PabA/PabB generic interface** |  |  |  |  |  |  |
| aY127A/wt | - | + | > 8 | 0.17 ± 0.02 | 0.02 ± 0.000 | 0.14 ± 0.01 |
| aS171A/wt | +/- | + | 0.92 ± 0.71 | 0.05 ± 0.00 | 0.08 ± 0.001 | 1.46 ± 0.08 |
| [Y127A, S171A]/wt | - | +/- | n. b. | 0.22 ± 0.01 | 0.01 ± 0.000 | 5.5 *10<sup>-2</sup> ± 4.0 *10<sup>-3</sup> |
| wt/bD308A | - | - | n. b. | 8.57 ± 1.29 | 0.11 ± 0.005 | 1.3 *10<sup>-2</sup> ± 2.5 *10<sup>-3</sup> |
| wt/bD308N | - | - | n. b. | 2.63 ± 0.30 | 0.10 ± 0.003 | 3.6 *10<sup>-2</sup> ± 5.5 *10<sup>-3</sup> |
| wt/bD308E | - | +/- | 1.70 ± 0.48 | 0.19 ± 0.01 | 0.29 ± 0.005 | 1.54 ± 0.12 |
| wt/bR311A | - | - | n. b. | 3.94 ± 0.44 | 0.39 ± 0.010 | 0.10 ± 0.01 |
| **B, PabA/PabB Gln-induced extended interface** |  |  |  |  |  |  |
| aR30E/wt | + | + | 0.38 ± 0.20 | 6.42 ± 0.58 | 0.51 ± 0.014 | 7.9 *10<sup>-2</sup> ± 9.3 *10<sup>-3</sup> |
| wt/bE119D | + | + | 0.84 ± 0.06 | 1.71 ± 0.05 | 0.59 ± 0.005 | 0.34 ± 0.01 |
| wt/bE119R | +/- | +/- | 0.54 ± 0.04 | 8.07 ± 0.83 | 0.48 ± 0.014 | 6.0 *10<sup>-2</sup> ± 7.9*10<sup>-3</sup> |
| **C, PabA/PabB glutaminase active site** |  |  |  |  |  |  |
| aP53A/wt | + | + | 0.18 ± 0.07 | 0.65 ± 0.02 | 0.41 ± 0.004 | 0.63 ± 0.03 |
| aC54G/wt | + | + | 0.06 ± 0.02 | 0.33 ± 0.02 | 0.43 ± 0.007 | 1.28 ± 0.11 |
| aC54A/wt | + | + | 0.18 ± 0.12 | 0.36 ± 0.04 | 0.45 ± 0.011 | 1.24 ± 0.16 |
| aH128A/wt | +/- | +/- | 0.37 ± 0.07 | 29.8 ± 1.73 | 0.28 ± 0.006 | 9.2 *10<sup>-3</sup> ± 7.5 *10<sup>-4</sup> |
| wt/bG207P | +/- | +/- | 1.70 ± 0.60 | 18.7 ± 1.01 | 0.11 ± 0.002 | 5.7 *10<sup>-3</sup> ± 4.3 *10<sup>-4</sup> |
| wt/bY210A | - | +/- | $5.50 \pm 2.90$ | $2.73 \pm 0.31$ | $0.80 \pm 0.028$ | $0.29 \pm 0.04$ |
| wt/bY210F | + | + | $0.53 \pm 0.10$ | $1.48 \pm 0.08$ | $0.75 \pm 0.011$ | $0.51 \pm 0.03$ |

<sup>a</sup> ADCS complexation deduced from retention volumes as determined by analytical SEC. Due to the limited stability and availability of Cho, all SEC measurements were performed in the absence of Cho. +/-, partial ADCS complex formation (*cf.* **Figure 5**).
<sup>b</sup> Reported mean and standard errors for $K_D$ values were calculated from two technical ITC replicates. n. d., no binding detected. To prevent the emergence of background signal originating from substrate turnover, all ITC measurements were performed in the absence of Cho and Gln.
<sup>c</sup> Activity measurements were performed in the presence of Cho, unless otherwise stated. Reported mean values and standard errors for the steady-state kinetic constants were calculated from a biological duplicate, each assayed in the form of at least two technical replicates.

Next, we determined the crystal structures of the ADCS complex with a number of reaction ligands, to provide insights into the conformational changes of the glutaminase active site environment prior to and during catalysis **(Figures 1, 2, Supplementary Figure 1a)**. All crystals used for structure determination contained two heterodimeric ADCS complexes with variable ligand occupancy **(Supplementary Table 1, Supplementary Figure 2)**. When Gln was added to the crystallization buffer, we found a glutamyl thioester (Gln-TE) adduct covalently bound to the catalytic aC79 within the glutaminase active site in only one of the two PabA/PabB heterodimers, indicating that under the given crystallization conditions the glutaminase reaction proceeded only to the Gln-TE adduct without performing the final hydrolysis step **(Figure 2, Supplementary Figure 1a)**. When Cho was subsequently added to these crystals, it was bound to the synthase active site of both ADCS heterodimers **(Figure 1, Supplementary Figure 3a-b, Supplementary Table 1)**. Consistent with previous observations for related MST enzymes ^15^, the benzoate moiety of Cho is complexed by a magnesium ion **(Supplementary Figure 4)**.

**Figure 1.**
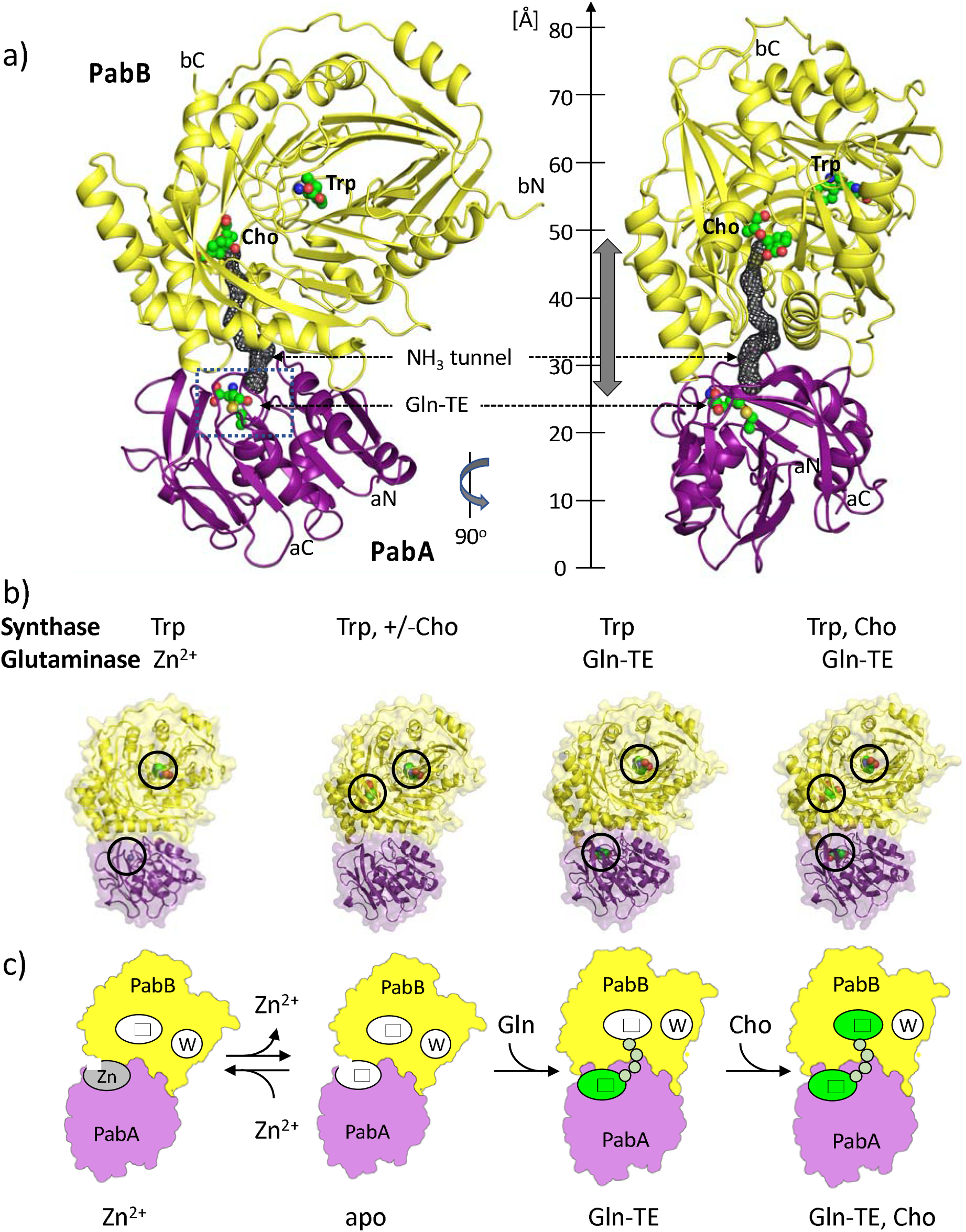
Overall architecture of the ADCS complex from *E. coli*. **a,** Cartoon representation of the ADCS complex formed by the PabA (violet) and PabB (yellow) subunits in two different orientations, rotated vertically by 90 degrees. The glutaminase and synthase active sites are marked by bound Gln-TE and Cho, respectively, in a combined spheres and sticks presentation (green, oxygen atoms in red, nitrogen atoms in blue, sulfur atoms in beige). The PabB Trp binding site and the N- and C-termini of both subunits are also labeled. The ammonia tunnel connecting the two active sites is shown by a grey mesh and its approximate length is indicated by a grey double arrow. The overall dimensions of the ADCS complex are indicated by a vertical ruler; **b,** Semitransparent surface/cartoon representations of ADCS structures with different glutaminase and synthase ligands bound: Trp, Zn^2+^ (PDB entry: 8RP6); Trp, +/-Cho (PDB entries: 8RP2, 8RP1 chains B/C, 8RP0 chains B/C); Trp, Gln-TE (PDB entry: 8RP1 chains A/D); Trp, Gln-TE, Cho (PDB entry: 8RP0 chains A/D). Bound ligands are shown in the respective structures and highlighted by circles; **c,** Schematic representation of ADCS structures in four catalysis states, designated Zn^2+^, apo, Gln-TE, and Gln-TE, Cho (*cf*. **Supplementary Figure 1a**). Note that there is no difference in the glutaminase active site in the presence of Gln-TE and of Gln-TE, Cho. The glutaminase and synthase active sites are marked both with asterisks. Zinc binding to the glutaminase active site is labeled “Zn”, Trp binding to PabB subunit is labeled “W”. Active sites loaded with substrate (Cho) or reaction ligands (Gln-TE) are colored in green. The protein tunnel connecting the glutaminase/synthase active sites is indicated by an array of small circles in light green.

**Figure 2.**
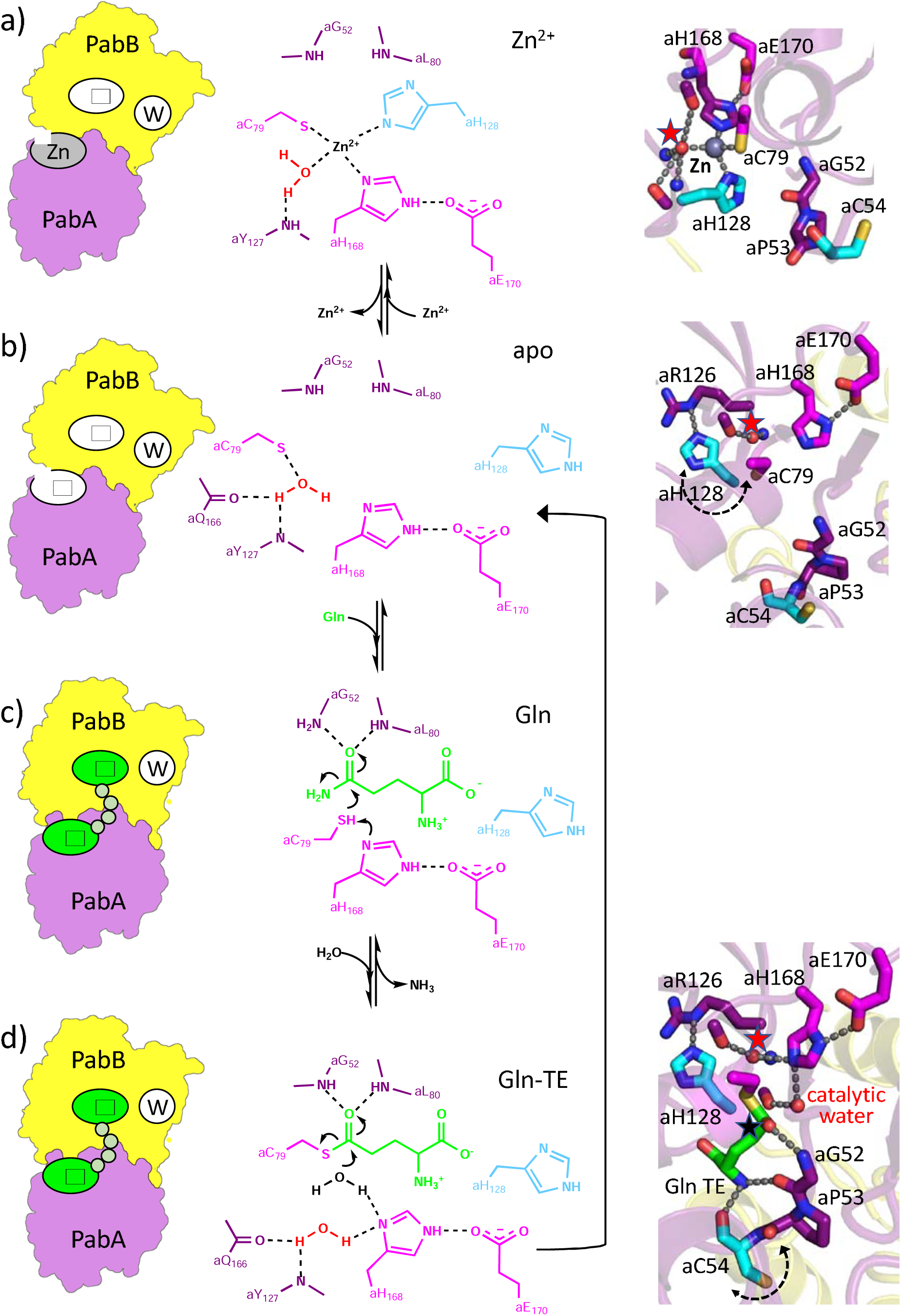
Structurally defined ADCS catalysis states. Left, schematic representations of ADCS structures (***cf*.** Figure 1c); middle, chemical representations of the glutaminase active site with expected electron movements during individual catalysis steps indicated by curved arrows; right, structural representations of the glutaminase active site. **a,** ADCS with Zn^2+^ bound into the glutaminase active site (PDB entry; 8RP6); **b,** ADCS (apo) without Zn^2+^ bound into the glutaminase active site (PDB entry: 8RP2); **c,** ADCS glutaminase active site upon Gln binding, indicating a catalytically competent state, to proceed to Gln-TE formation **(*cf*. Supplementary Figure 6); d,** ADCS glutaminase active site with bound Gln-TE, indicating a catalytically competent state to allow Gln-TE hydrolysis (PDB entry: 8RP1 chains A/D). The mobile glutaminase active site residues aC54 and aH128 are in cyan, and the approximate mobility directions are indicated by dashed double arrows. the glutaminase catalytic triad residues aC79, aH168 and aE170 are in pink; glutaminase active site oxyanion residues aG52 and aL80 are in deep purple. Glutaminase reaction ligands are in green. Hydrogen bonds are shown by dashed lines. In the structural representation, the reactive carbon atom on Gln-TE, subject to nucleophilic ligand substitution, is indicated by a black asterisk. The structurally conserved *glutaminase active site-forming water molecule* is marked by a red asterisk. For the sake of clarity, the orientation of the middle and right panels differs by approximately 90 degrees around an axis perpendicular to the viewing plane, and only a subset of active site residues is shown in the chemical representations (middle panel).

The overall size of the ADCS assembly exceeds 80 Å in its longest dimension and its subunit arrangement is related to structures of other members of the MST family that use ammonia as nucleophile **(Figure 1a, Supplementary Figure 1b, Supplementary Table 2)**. In all ADCS structures, we found a structurally conserved Trp molecule bound to each PabB synthase subunit **(Figure 1, Supplementary Figure 3c, Supplementary Table 1)**, in agreement with previous structural data of closely related synthase subunits or domains of ADCS and anthranilate synthase (AS) ^15^. Contrary to its role as product inhibitor in the AS complex, we could not detect any inhibiting effect of Trp in ADCS activity, in agreement with previously published biochemical data ^21,22^.

We also identified a tunnel connecting the two active sites, which is required for the glutaminase product ammonia to be transferred to the synthase active site **(Figure 1a, Supplementary Figure 4)**. Unlike observations in other GAT structures, parts of this tunnel appear to be too narrow to allow unhindered ammonia transport ^2–4^, thus requiring conformational adaption during ADCS catalysis. The tunnel is completely sequestered from external solvent and is about 27 Å in length. The entrance to the tunnel is adjacent to the glutaminase active site at the PabA/PabB interface and shared by residues from both subunits. The central part of the tunnel is mainly formed by hydrophobic PabB residues and terminates next to the 4-hydroxy-1,5-cyclohexadiene-1-carboxylic acid ring of Cho, which is the reactive group for Cho transamination ^15,16^.

### ADCS comprises a structural zinc binding site next to the glutaminase active site

Interestingly, in the ADCS structure without any reaction ligands bound to the glutaminase active site, we observed a metal ion with tetrahedrally coordinated interactions to the two PabA catalytic triad residues aC79 and aH168 as well as to aH128 next to the PabA active site, as evidenced by an anomalous electron density map **(Figures 1b-c, 2a, Supplementary Figure 3e)**. As the coordination geometry of this site did not match any of the ions added during ADCS purification or crystallization (Na^+^, K+, Mg^2+^), we searched for alternative metal ions by X-ray fluorescence (XRF) analysis (**Supplementary Figure 5a)**. The resulting data allowed assigning the bound metal ion as Zn^2+^ that fits well with the predicted coordination geometry ^23,24^. The presence of Zn^2+^ was independently confirmed by XRF of ADCS in solution **(Supplementary Figure 5b)**. In contrast, we did not detect any Zn^2+^ in an ADCS variant, in which one of the coordinating ligands was removed (aH128A), demonstrating that it originates from the structurally observed Zn^2+^ site **(Supplementary Figure 5c)**. When the crystallization buffer was incubated with EDTA, in the resulting X-ray structure the density assigned to Zn^2+^ also disappeared **(Figures 1b-c, 2b, Supplementary Figure 5c)**. In ADCS complex structures obtained in the presence of Gln, we did not observe any Zn^2+^ binding to the glutaminase active site **(Supplementary Table 3)**. As Zn^2+^ was not added at any stage during purification and crystallization, the most plausible explanation for its presence is by its extraction from the expression host.

Comparison of the glutaminase active site geometry in the presence and absence of Zn^2+^ allowed to identify conformational changes associated with Zn^2+^ binding **(Figures 2a-b)**. While we did not detect any significant positional changes for the catalytic triad residues aC79, aH168 and aE170, the side chain of aH128 flips from an orientation blocking the PabA active site in the presence of Zn^2+^ to a conformation, in which it integrates into an interaction network forming the glutaminase active site in a catalytically competent conformation, in the absence of Zn^2+^. However, our biochemical data did not reveal any effect of Zn^2+^ on ADCS activity (Supplementary Table 4).

### The ADCS glutaminase active site is shared by the PabA and PabB subunits

Next, we investigated the structural requirements to allow the formation of a complete and catalytically competent ADCS glutaminase active site. When Gln forms a covalent adduct with the catalytic aC79, as expected the glutaminase catalytic triad residue aH168 is in a catalytically competent position, allowing a nucleophilic attack of water on the reactive carbon atom to reconstitute free aC79 **(Figures 2d, 3a, Supplementary Figure 1a)**. In the ADCS structures with Gln-TE bound in the glutaminase active site, we consistently observed a distinct water molecule interacting with the active site residues aY127 and aH168. This water molecule is in an appropriate position to hydrolyze the Gln-TE, as illustrated by a close distance of less than 3 Å to the Gln-TE reactive carbon atom, which is subject for nucleophilic substitution, and hence it is referred to as the *glutaminase catalytic water molecule*. Finally, the reactive carbonyl group of the Gln-TE interacts with the main chain amino groups of aG52 and aL80 **(Figures 2c-d, 3a)**, which together form the oxyanion hole required for glutaminase catalysis by polarization of its carbonyl group. Contrary to findings in other GATs ^11,25–27^, these residues do not undergo significant conformational changes upon Gln binding, except for aC54 adjacent to aG52 (see below).

We also found several interactions between the non-catalytic amino group of the Gln-TE and residues of both ADCS subunits, including bG207 and bY210 from the PabB subunit **(Figure 3a)**, allowing the catalytic center of Gln-TE to be positioned with very high spatial precision, to generate a catalytically competent glutaminase active site arrangement at the PabA/PabB interface. In this arrangement, the entire exposed Gln-TE surface not interacting with PabA residues (25 Å^2^) is buried by interactions with residues of the PabB subunit, demonstrating that the ADCS glutaminase active site is shared with the synthase subunit PabB. Finally, in all ADCS complex structures without Zn^2+^, we found a second well-defined and structurally conserved water molecule in the glutaminase active site **(Figure 2)**. It is located approximately midway between the imidazole side chain groups of aH128 and aH168, suggesting that it critically contributes to establish a catalytically competent conformation of the ADCS glutaminase active site.

**Figure 3.**
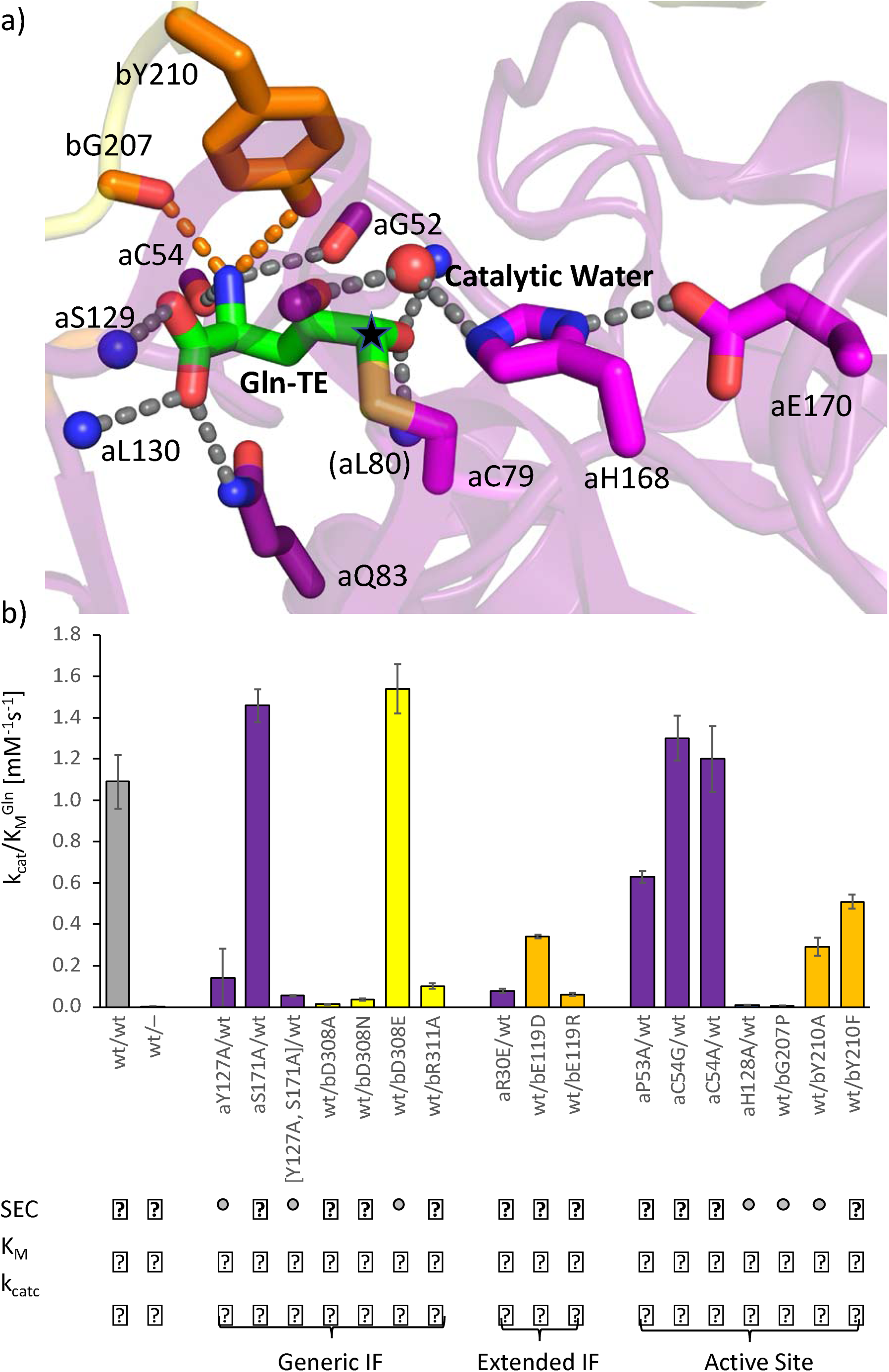
Effect of PabA/PabB interface formation on a catalytically competent glutaminase active site structure. **a,** Gln-TE bound to the glutaminase active site in a catalytically competent conformation. Hydrogen bonds are indicated by dashed lines. The Gln-TE carbon atom, where the substitution of the substrate amino group leading to TE formation and subsequent Gln-TE hydrolysis takes place, is indicated by a black asterisk. **b,** Histogram of catalytic efficiencies k_cat_/K_M_ of biochemically and biophysically characterized ADCS variants (***cf*. Table 1**). Estimated standard deviations from repeated measurements (N=4) are shown by errors bars. ADCS variants are grouped in mutants affecting the generic Gln-independent interface (IF), the Gln-TE-induced extended interface, and the glutaminase active site. Note that the residues listed under the active site category are also part of the extended PabA/PabB interface **(*cf*.** Figure 4). The color codes are as in previous figures. PabB residues that contribute to the extended interface are in orange. Effects on ADCS complex formation estimated by SEC analysis are indicated below: little or no effect on ADCS complex formation, black circles; major effects on ADCS complex formation, grey circles; no ADCS complex formation, white circles. Effects on the Michaelis constant K_M_ and the catalytic constant k_cat_ are also indicated; > 10 % of the value of *wt* ADCS, black circles; < 10% of the value of *wt* ADCS, white circles. In several ADCS variants, the diminishment of ADCS complex formation and the Gln binding affinity, estimated by K_M_, are correlated.

We also produced a model of ADCS with the glutaminase substrate Gln bound to its active site, using an energy minimization protocol by using ADCS with bound Gln-TE as template **(Figure 3a, Supplementary Figure 6a)**. It illustrates that Gln is identically bound to the glutaminase active site. The modeled Gln position was further confirmed by structures of other GATs with class I glutaminases in complex with Gln ^5^ **(Supplementary Figure 6b)**. Based on these findings, we use our experimentally determined ADCS Gln-TE complex structures as a template for the positioning the glutaminase substrate (Gln) in ADCS elsewhere, especially for the interpretation of our biochemical and biophysical data see below).

### Glutamine binding induces sequestration of the ADCS glutaminase active site through PabA/PabB subunit interface expansion

Next, we investigated whether the presence of ADCS reaction ligands (Gln-TE, Cho) leads to changes in the overall PabA/PabB subunit arrangement. Indeed, we found a rotation by approximately 23 degrees in the principal axis orientation of PabA and PabB subunits induced by the presence of Gln-TE (**Figure 4a-b, Supplementary Figure 7, Supplementary Table 3)**. In contrast, upon additional Cho and Mg^2+^ binding to the ADCS synthase active site, there were no further significant alterations in the PabA/PabB arrangement, consistent with only a minor regulatory effect of Cho/Mg^2+^ on glutaminase activity **(Supplementary Table 4)**. When analyzing how these changes in PabA/PabB arrangement affect the interface between the two subunits, we found a 30% increase from an area covering the generic, substrate-independent PabA/PabB interface of slightly less than 1000 Å^2^ to approximately 1300 Å^2^ in the presence of Gln-TE **(Supplementary Table 3)**. This interface expansion leads to complete sealing of the PabA active site rendering Gln-TE inaccessible to solvent, which is a prerequisite for sequestered ammonia transfer to the PabB active site **(Figure 4c-d, Supplementary Figure 4)**.

**Figure 4.**
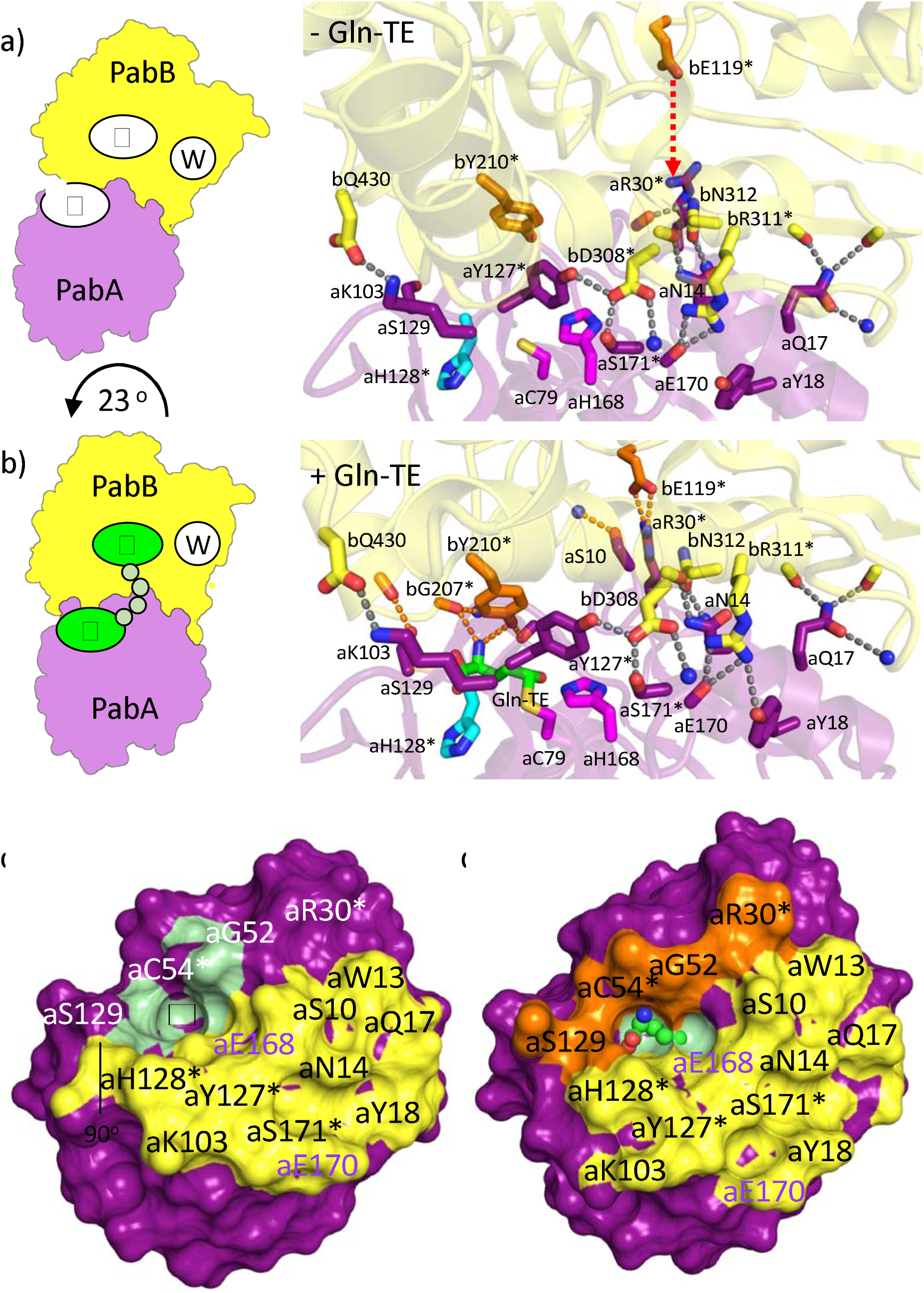
ADCS PabA/PabB interface changes induced by Gln-TE. **a,** Structural representation of the PabA/PabB interface with no ligands bound (PDB entry 8RP1 chains B/C) and **b,** in presence of Gln-TE (PDB entry 8RP1 chains A/D). When Gln-TE is bound to the glutaminase active site, the PabA/PabB interface is expanded by about 1/3 of the area of the generic, Gln-independent interface. Residues contributing to the extended interface are shown in orange. The movement of bE119 to form a salt bridge with aR30 in the presence of Gln-TE is indicated by an arrowed line in red, caused by a 23° rigid-body rotation of the PabA/PabB arrangement **(Supplementary Figure 7).** Corresponding ADCS PabA/PabB arrangements are shown on the left, as defined in previous figures. Colors are as in previous figures. PabB residues and hydrogen bonds to PabA residues of the extended interface are in orange. Residues that have been biochemically and biophysically characterized **(Table 1)** are marked with asterisks. **c,** Generic PabA surface interacting with PabB in the absence of Gln-TE. **d**, PabA Gln-TE induced extended surface interacting with PabB. The glutaminase active site without Gln-TE is indicated by a black asterisk; Gln-TE is shown by ball/stick representation. Only when Gln-TE is present, the glutaminase active site is completely wrapped by residues contributing to the generic (yellow) and extended (orange) interface with PabB. The remaining glutaminase active site surface is in pale green.

While the PabA/PabB interfacial interactions observed in the absence of Gln-TE remained in its presence, we consistently found a number of additional direct PabA/PabB interfacial interactions that are present only in ADCS complexes with bound Gln-TE **(Figures 3a, 4a-b, Supplementary Table 5)**. The associated changes are most visibly illustrated by a long-range movement of residue bE119 from the PabB subunit, which is 10-13 Å away from any PabA residues in different ADCS structures devoid of Gln-TE in the glutaminase active site. In contrast, when Gln-TE is present, bE119 interacts via a direct salt bridge with aR30 from the PabA subunit, remote from the glutaminase active site. This interaction allows additional glutaminase residues adjacent to or directly contributing to the glutaminase active site, to become part of the extended PabA/PabB interface and thus completely enclosing the active site. In particular, a PabA sequence segment (aG52-aP56), which includes a part of the oxyanion hole of the glutaminase active site, undergoes significant positional movements only upon Gln-TE binding **(Figure 2, Supplementary Figure 8a)**. By analyzing the dihedral angles of the peptide bonds of this segment, we found major changes for residues aP53 and aC54, inducing a highly strained conformation of aC54 in the presence of the Gln-TE **(Supplementary Figures 9, 10)**.

### ADCS glutaminase activity requires PabA/PabB subunit interface expansion

To investigate the functional impact of the PabA/PabB interface expansion during glutaminase catalysis, we analyzed the ADCS variant complexes by analytical size exclusion chromatography (SEC) in the presence and absence of Gln **(Figures 3b, 5, Table 1)**. A significant proportion of these variant complexes showed Gln-dependent elution profiles, demonstrating the feasibility of this approach to evaluate the effects of Gln on enhanced PabA/PabB complex formation. We also quantitatively measured the binding affinity of PabA/PabB variant complexes in the absence of Gln by ITC.

First, we focused on two PabB residues bD308 and bR311, which represent highly conserved PabB sequence positions and critically contribute to the generic PabA/PabB interface of the ADCS complex, independent of the presence of glutaminase substrate ^21,28,29^. Due to their exposed PabB surface positions their side chains extend into distinct PabA surface pockets, which are centrally situated in the overall PabA/PabB interface **(Figure 4)**. We observed that side chain replacement by alanine in each of them is sufficient to cause complete PabA/PabB disassembly, even in the presence of Gln **(Figures 3b, 5c, Table 1, section A)**. To further test the specific role of bD308, we also mutated this residue into an isosteric mutant bD308N without a negative charge and to bD308E, which is extended by one methylene group. As with the bD308A variant, we did not observe any PabA/PabB complex formation in the bD308N variant also in the presence of Gln, demonstrating that the side chain carboxylate group is essential for PabA/PabB assembly. When testing the bD308E variant, PabA/PabB assembly was partially restored when Gln was added. Except for the bD308E mutant, we found only residual glutaminase activity in all other bD308 and bR311 mutants. Consistent with the observed loss of PabA/PabB complex formation, the increased K_M_ ^Gln^ values of these mutants indicated substantially weakened Gln binding **(Table 1)**. As these PabB residues are not directly involved in the formation of the glutaminase active site, our findings demonstrate that stable PabA/PabB complex formation in the presence of Gln is essential for glutaminase activity.

When changing single PabA residues aY127 and aS171 or a combination of both to alanine, both interacting with bD308, no or only a weak tendency for complex formation was observed in the absence of Gln as well. However, the addition of Gln restored ADCS complex formation **(Figure 5d**, **Table 1)**. The mutants with a partial PabA/PabB disassembly effect also retained a high level of substrate binding. In contrast, glutaminase turnover number in these variants was much more affected than in ADCS variants, in which interacting PabB residues bD308 and bR311 were mutated. These data demonstrate a direct involvement of aY127 and aS171 in glutaminase catalysis, which can be explained by their sequential and structural proximity to the catalytic residues aH168 and aE170.

**Figure 5.**
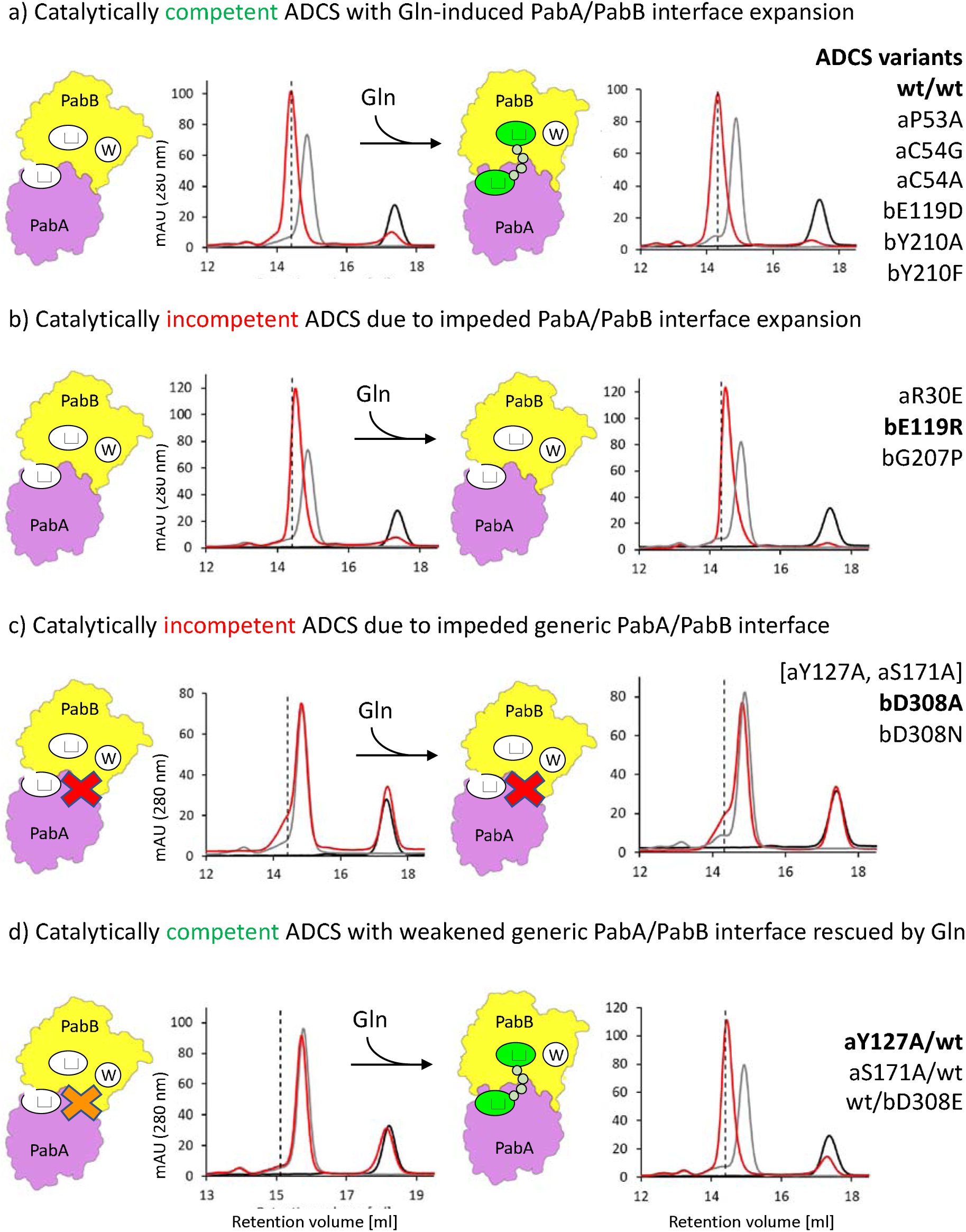
Contributions of PabA/PabB interface residues to ADCS complexation and glutaminase activity. Representative SEC profiles in red indicate the ADCS assembly state before (left panels) and after (right panels) adding Gln. The SEC profiles of separate PabA and PabB subunits, eluting at approximately 17.4 and 14.9 mL, respectively, are shown in black and grey. Fully assembled ADCS complex elutes at approximately 14.4 mL, as indicated by the vertical dashed lines. ADCS variants that fall into one of the four categories (***cf*. Table 1**) are listed on the right. ADCS variants, for which representative data are shown, are in bold: **a**, Gln-induced PabA/PabB interface expansion generating a catalytically competent ADCS complex; **b**, ADCS variants impeded for PabA/PabB interface expansion, causing diminishment of ADCS glutaminase activity to residual levels; **c**, ADCS variants with impeded generic PabA/PabB interface (indicated by red cross), which cannot be rescued by Gln addition and thus have only residual catalytic activity**; d,** ADCS variants with weakened generic PabA/PabB interface (indicated by orange cross) where complexation is rescued by Gln-induced interface expansion to generate partially active ADCS complexes. The ADCS assembly and catalysis states are also indicated schematically (***cf*.** Figure 1c). The complete set of SEC profiles is shown in **Supplementary Figure 14**.

To investigate the molecular basis of interface expansion for ADCS glutaminase catalysis, we introduced charge-reversing side chain substitutions in both residues of the aR30-bE119 salt bridge, which is main contributor of the Gln-TE induced extended PabA/PabB interface, remote from the glutaminase active site. As expected, both ADCS aR30E and the bE119R variants had only a minor effect on PabA/PabB complex formation in the presence and absence of Gln but showed only residual glutaminase activity due to substantially weakened Gln-binding capacity during catalysis **(Figures 3b, 5b, Table 1)**. These data demonstrate the crucial role of the aR30-bE119 salt bridge in the formation of an extended PabA/PabB interface.

Next, we focused on possible contributions of residues from the extended interface of both ADCS subunits directly involved in Gln-TE binding for ADCS glutaminase activity, starting with PabB residues bG207 and bY210 **(Figures 2a-b, 3a, 4)**. Judging from our structural data these PabB residues, participating in the glutaminase active site, are too distant to participate in Gln binding when interface expansion is impeded. In addition, bG207 adopts a dihedral angle conformation in all PabA/PabB complex structures of this contribution, which would be non-permissive for non-glycine residues, explaining its high level of conservation ^20^. Using an ADCS variant with bG207 mutated to proline (bG207P) leads to compromised PabA/PabB complex formation, both in the absence and in the presence of Gln **(Figure 5b**, **Table 1)**. Correlated with this, we found a 55-fold diminishment of substrate binding affinity, reducing substrate turnover to residual levels. These data demonstrate a crucial role of this PabB residue to complete the ADCS glutaminase active site and to contribute to catalytic activity. Side chain removal of bY210 (bY210A) induces an even stronger PabA/PabB assembly defect but the effects on glutaminase activity are less severe, demonstrating a less crucial involvement in glutamine binding. In contrast, biochemical analysis of PabA variants of residues aP53 and aC54 revealed only minor effects on glutaminase activity **(Table 1)**, which is plausible as they remain close to the glutaminase active site irrespective of interface expansion.

Finally, we assessed whether a catalytically competent conformation of the glutaminase active site requires rigidification by the presence of the synthase subunit PabB, by determining an additional crystal structure of the separate PabA subunit **(Supplementary Figure 11, Supplementary Table 1)**. Indeed, comparison of these structures reveals a significant increase in ordering of the glutaminase active site in the presence of PabB due to reduced conformational variability of several interface PabA loops, which is required to generate a catalytically competent arrangement of the two catalytic triad residues aH168 and aE170. Along these lines, we also investigated the role of the mobile residue aH128 next to the glutaminase active site by removing its side chain **(Figures 2, 4)**. The residual glutaminase activity observed for the aH128A variant is due to an approximately 100-fold increase in K_M_, which reflects an almost loss of Gln-binding capacity **(Table 1)**. As this variant showed only a minor effect on PabA/PabB assembly, the imidazole side chain of aH128 appears to play a crucial role in glutamine binding by contributing to a catalytically competent glutaminase active site conformation.

In summary, our structural and biochemical data of the ADCS complex reveal separate requirements for the initial formation of a glutaminase substrate-independent generic and an additional extended PabA/PabB interface upon glutaminase substrate (Gln) binding, mediated by a rearrangement of the two subunits by a 23 degrees rotation **(Figure 5)**. Both interface segments have areas remote from and contributing to the glutaminase active site, but when impeded, the effects on ADCS assembly and activity are different. Depending on whether impairment of the generic PabA/PabB interface leads to reduced ADCS assembly, partial ADCS glutaminase activity can be rescued, but not in ADCS variants where complete disassembly was observed **(Figure 5c-d)**. In contrast impairment of the extended PabA/PabB interface has little or no effect on ADCS assembly, as the generic substrate-independent assembly still remains, but it results in catalytically inactive ADCS, as a catalytically competent glutaminase active site cannot be formed and its sequestration is lost **(Figure 5b)**. For residues that contribute to the extended PabA/PabB interface but are not essential for its formation, the effects are stronger for PabB variants that fail to properly position into the ADCS glutaminase active site shared by both subunits than for PabA variants that remain close to the glutaminase active site even when the PabA/PabB interface is impeded **(Figure 5a).**

## DISCUSSION

Evolution has optimized multienzyme systems to be available at very precise stoichiometries, which are associated with repair mechanisms when these compositions become unbalanced ^30–32^. In contrast, only little is known about the mechanisms by which corresponding multiple catalytic activities are regulated to avoid unbalanced and potentially wasteful consumption of individual metabolites. Contrary to unrelated glutaminases that are broadly involved in ammonia homeostasis and metabolism ^33,34^, ammonia generated by GAT-catalyzed Gln hydrolysis, needs to be tightly sequestered from external solvent during its transport to the synthase active site to remain in the reactive unprotonated state. Therefore, there is a strict functional requirement for closing the glutaminase active site during glutaminase catalysis and to properly position ammonia for subsequent transport to the synthase active site. As GAT class I glutaminases belong to the α/β hydrolase fold with numerous examples of active enzymes without a need for complex formation with other enzyme partners ^35,36^, additional mechanisms have evolved to regulate metabolite homeostasis and to avoid wasteful metabolite consumption ^7,25^.

In this contribution, using ADCS as model, we have demonstrated a mechanism in which GAT glutaminase activity is controlled by glutaminase active site sequestration mediated by glutamine-induced protein/protein interface expansion. These effects are generated by a 23 degrees rigid body rotation of the two interacting subunits PabA and PabB (Figure 4, Supplementary Figure 7, Supplementary Table 3). Consistent with these observations, we have shown that a weakened generic, PabA/PabB interface can be rescued upon incubation with Gln, allowing recovery of catalytical competence **(Figure 5d)**. In contrast, variants with perturbed interface expansion are impaired in their ability to bind and hydrolyze Gln **(Figure 5b)**.

When searching for similar effects to generate GAT arrangements in other members of the MST family, we indeed detected interface expansion in those systems, where a comparison of glutaminase states in the absence and presence of Gln or subsequent catalysis intermediates was possible ^37,38^. However, in none of these related GATs, could we detect a significant rigid body movement of the two participating subunits (or domains in the case of fusion proteins) at the scale observed for ADCS. Whether this is due to a lack of suitable experimental conditions in these systems investigated or whether such a regulatory mechanism does not exist, remains currently unknown. Interestingly, we found one distantly related GAT system, imidazole glycerol phosphate synthase (IGPS, HisH/HisF), in which the two subunits likewise rearrange by a rotation of > 20 degrees upon addition of Gln ^8,25^. Remarkably, this system additionally requires the presence of synthase substrate bound to the synthase active site, to activate glutaminase catalysis by a long-distance allosteric mechanism, which converts the glutaminase active site oxyanion hole to a catalytically competent conformation. In contrast, in ADCS the glutaminase active site oxyanion hole is in an active conformation regardless the addition of any substrates and there is no significant allosteric activation mediated by substrate binding to the ADCS synthase active site **(Figure 2, Supplementary Table 4)**.

Since there is no evidence for allosteric synthase substrate-induced glutaminase activation in ADCS at a scale comparable to the IGPS system ^8,12^, we wondered about any additional mechanisms of glutaminase activity regulation in ADCS and related GAT members of the MST family **(Supplementary Table 6)**. Interestingly, in the related ADICS complex a Zn^2+^ binding site was found, which is in an almost identical position as observed in our ADCS data ^37^ **(Supplementary Figure 13)**. Whether such site is preserved in other GAT members of the MST family remains currently unknown. As the activities of several enzymes involved in folate biosynthesis are either directly controlled by the presence of Zn^2+^ or their expression is under the control of zinc-dependent promoters such as the Zur promoter, Zn^2+^ binding in ADCS may contribute to Zn-mediated homeostatic regulation ^39–41^. Whether, for instance, liberation of zinc during ADCS catalysis has any effect on the pool of available zinc remains an attractive hypothesis, given that the availability of free zinc in living organisms is negligible ^42,43^.

Considering the additional effects detected with moderate potential of ADCS glutaminase activity regulation, our data demonstrate that GAT glutaminase/synthase surface expansion together with major rearrangements of the participating subunits (or domains in case of fused enzyme versions) is likely to be a pivotal point of overall GAT activity regulation. Their quantitative structural and biochemical characterization in other GATs remains a fascinating topic for future research, to unravel general principles in GAT activity synchronization, ultimately allowing us to learn about naturally evolved mechanisms to ensure the most efficient utilization of metabolites in catalytic processes.

## METHODS

### Cloning and Mutagenesis

The genes for ecPabA and ecPabB were cloned into a pET28a vector with an N-terminal His_6_-Tag followed by a TEV cleavage site ^44^. To increase the efficiency of TEV cleavage a G5H-Tag was introduced between the TEV cleavage site and the methionine start codon of PabA via site directed mutagenesis, which concomitantly improved solubility and stability of PabA. PabA and PabB variants were also produced by site-directed mutagenesis.

### Gene Expression and Protein Purification

*E. coli* BL21-Gold (DE3) cells were transformed with the respective plasmid and grown in 3-6 L Luria broth medium supplemented with 50 µg/ml kanamycin at 37 °C to an OD_600_ of 0.6. Expression was induced by 0.5 mM IPTG, followed by overnight incubation at 25 °C ^20^. Cells were harvested, resuspended in 50 mM Tris/HCl pH 7.5, 300 mM KCl, 10 mM imidazole and 2 mM DTT, and lysed by sonication.

Proteins were purified via Ni-immobilized metal affinity chromatography (IMAC) by applying the soluble lysate to a HisTrap FFcrude column (5 ml, GE Healthcare), which was developed with a linear gradient of imidazole (0-500 mM). Proteins used for crystallization were cleaved with TEV protease (TEV protease to protein mass ratio: 1 to 50) while dialyzing against 3 L 50 mM Tris/HCl pH 7.5, 50 mM KCl at 4 °C overnight. After TEV cleavage, TEV protease, the split-off His-Tag and uncleaved protein was removed by reverse IMAC. Fractions containing the proteins of interest were identified by SDS-PAGE, pooled and further purified by preparative gel filtration on a Superdex75 pG HiLoad 26/600 column (320 ml, GE Healthcare). Protein concentrations were determined spectrophotometrically at 280 nm, with molar extinction coefficients calculated via ExPASy ProtParam.

### Cho Purification

Chorismate (Cho) was purified from culture supernatant of *E. coli* KA12 cells as described previously ^45^. After removing the cells by centrifugation, the supernatant was acidified with HCl (5 M) to pH 1.5 and extracted with ethylacetate (2 x 180 ml). The combined extracts were washed with saturated brine (360 g/l NaCl) and ethylacetate was removed *in vacuo*, yielding a small volume of crude Cho. The material was loaded on 50 g C18 reverse phase silica and eluted with 10 mM ammoniumacetate (pH 6.0), 10 ml fractions were collected and for each fraction absorption spectra were recorded. Peak fractions containing Cho (ε_275_(Cho) = 2630 M^-1^ cm^-1^) were combined and lyophilized. The resulting lyophilizate was dissolved in a small volume of sterile water, aliquoted, and stored at −80°C. Sample purity was assessed by analytical reverse phase HPLC and total turnover by anthranilate synthase in an enzymatic assay ^44^.

### Analysis of ADCS complex formation by analytical SEC

Analytical SEC was applied to assess the ability of subunit variants to form a heterodimeric ADCS complex in the presence or absence of Gln ^20^. PabA and PabB subunits were mixed in an equimolar ratio to final concentrations of 50 µM each in SEC buffer (50 mM Tris/HCl pH 7.5, 50 mM KCl, 5 mM MgCl_2_, 2 mM DTT and, where indicated, 5 mM L-Gln). As control, the isolated *wt* subunits were also applied at 50 µM concentration under the same measuring conditions. Samples were applied to a Superdex200 increase 10/300 GL column (GE Healthcare) equilibrated with SEC buffer with or without L-Gln at 25 °C at a flow rate of 0.3 ml/min. Protein elution was followed by absorbance measurements at 280 nm. Calibration was performed with the Cytiva LMW and HMW calibration kits.

### Analysis of ADCS complex formation by ITC

ITC measurements were performed to quantify ADCS complex affinities. PabA and PabB subunits were dialyzed (Spectra/Por® 1 Dialysis Membrane, MWCO 6-8 kDa) overnight at 4 °C against a 1000-fold volume excess of degassed ITC buffer (50 mM Tris/HCl pH 7.5, 50 mM KCl, 5 mM MgCl_2_). Experiments were performed using a MicroCal PEAQ-ITC instrument (Malvern) at 25 °C with a reference power of 10 µcal/s. A total of 19 or 23 injections of 1 or 2 µl 250 µM PabB were titrated from a rotating syringe (750 rpm) into the isothermal chamber containing 300 µl 25 µM PabA with a delay time of 150 s between injections. Enthalpy differences were obtained by integrating the heat pulses of each injection and plotted against the PabB/PabA ratio. From the resulting isotherm, the binding stoichiometry N, dissociation constant K_D_ and thermodynamic parameters ΔH and ΔS were calculated using the MicroCal PEAQ-ITC Analysis Software (Fit: One set of sides, Version 1.41, Malvern).

### Steady-state enzyme kinetics

Glutaminase activity was measured under steady-state conditions by a continuous coupled enzymatic assay with glutamate dehydrogenase (GDH), coupling turnover of Gln to the production of α-ketoglutarate and the conversion of NAD^+^ to NADH (Δε_340 nm_ = 6220 M^-1^ cm^-1^) ^20^. Measurements were carried out in 50 mM Tricine pH 8, 5 mM MgCl_2_, 150 mM KCl, 1 mM DTT, 10 mM NAD^+^, 1 mg/ml GDH, 200 µM Cho and varying Gln concentrations. PabA variants were added at concentrations of 0.25 µM or 0.5 µM, PabB variants were added at a 4-fold excess over PabA. For measurements in the absence of PabB, the PabA concentration was increased to 5 µM. Measurements were conducted at 25 °C in 96-well plates using a plate reader (Tecan Infinite M200 Pro). To calculate glutaminase activity k_app_, the baseline-corrected initial slope of the reaction time course was transformed to 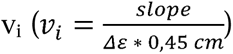 and normalized by the PabA concentration. To obtain the Michaelis-Menten parameters k_cat_ and K_M_, k_app_ was plotted against the concentration of Gln and a hyperbolic fit was applied to the resulting plot using the Origin 2020 software 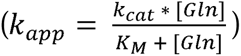. The reported values and standard deviations are based on a biological duplicate or triplicate, each with two technical replicates. Standard deviations of k_cat_/K_M_ values were calculated using Gaussian error propagation.

### Crystallization

ADCS crystals were grown at 18 °C in 24-well plates by combining 2 μl volumes of 8.25 mg/ml protein and reservoir solution at a 1:1 ratio, using the hanging-drop vapor diffusion method. The reservoir solution contained 0.1 M MES (pH 6.5) and 1.5 M magnesium sulfate heptahydrate. For ADCS structure determination in the presence of Gln-TE, the reservoir solution was supplemented with 0.2 M L-Gln After four days, crystals of mixed sizes were observed. Crystals larger than 50 μm in length were collected from the initial crystallization drop and washed with the reservoir solution five to six times. Subsequently, the washed crystals were crushed with a seed bead (JBS Beads-for-Seeds, CO-501) in 500 μl of precipitant and serially diluted 10, 100 and 1000 times. Macro-seeding was employed to obtain crystals of sufficient size for X-ray diffraction experiments and data collection. Crystals of a size of at least 300 x 200 x 100 μm^3^ were obtained after 3-4 days of macro-seeding using the hanging-drop vapor-diffusion method. For ADCS structure determination in the presence of Cho, 14.4 mM Cho was added to ADCS crystals co-crystallised in the presence of L-Gln. For ADCS structure determination in the absence of zinc, 1 mM EDTA was added to ADCS crystals without L-Gln added. All crystals were transferred into paratone-N oil as a cryoprotectant and flash-frozen in liquid nitrogen for storage, prior to X-ray data collection.

The separate PabA subunit was crystallized at 18 °C from 2 μl protein solution at 16.5 mg/ml and 2 μl reservoir solution comprising 0.2 M sodium formate, 20 % (v/w) PEG-3350, and 0.2 M L-Gln, employing the hanging drop vapor diffusion method. Small crystals were observed after one week. The cover glass was then transferred to 1 ml of new reservoir solutions with incrementally increased concentrations of 25%, 30%, and 35% (v/w) PEG-3350 every second day for sequential dehydration approaches. The dehydrated crystals were subsequently fast frozen in liquid nitrogen after equilibrating in 40% (v/w) PEG-3350 for two days.

### X-ray data collection, structure determination, and refinement

Diffraction datasets were collected at a wavelength of 0.9762 Å under cryogenic conditions (−180 °C) at the EMBL/DESY P13 beamline. XDS ^46^ was used for initial integration, followed by merging and scaling using the CCP4 suite program AIMLESS ^47,48^. Molecular replacement was conducted within CCP4i2 ^49^ using Phaser ^50^ with AS from *Serratia marcescens* as the search model (PDB code: 1I7Q) ^38^. Manual model building and refinement were performed using COOT ^51,52^ and REFMAC ^53^, respectively. The final refinement statistics are provided in Supplementary Table 1. The coordinates and structure factors have been deposited in the Protein data Bank (PDB) with accession codes 8RP0, 8RP1, 8RP2, 8RP6, 8RP7. The model of the Gln-bound ADCS complex **(Supplementary Figure 6a)** was generated by energy minimization using the YASARA force field and server (www.yasara.org/minimizationserver.htm) ^54^.

### Zinc ion identification

X-ray fluorescence (XRF) spectroscopy was utilised to identify metal ions present in the ADCS structure ^55^. All spectra were obtained using concentrated enzyme samples in 40% of glycerol and collected under cryo-conditions. The identification of zinc was based on the presence of the characteristic Kα emission energy peak at 8.6 keV. In all cases, the buffer solution used showed no trace of zinc. Using the anomalous differences from the ADCS dataset without any reaction ligands added and phases calculated with Refmac5 ^53^ from the apo ADCS structure (8RP6), anomalous difference maps were calculated using CCP4 suite ^47^. The maps were examined in Coot ^52^. In each of the PabA subunits within each asymmetric unit, we modelled a tetrahedral zinc site. To validate the observed geometry of these sites, we analyzed them using the CheckMyMetal server (CMM; http://www.csgid.org/csgid/metal_sites) ^24^.

### Sequence, structure and chemistry analysis and visualization

The sequence of PabA from *E. coli* (P00903) was used as template for a BLAST search using Uniref90 as database (www.uniprot.org/blast). Representative sequences of 27 sequence clusters, following the conditions to comprise at least ten different sequences and an unambiguous identification as glutaminase sequences with PabB as the only partner for complex formation, were submitted to Clustal Omega for multisequence alignment ^56^. The alignment was submitted to the WebLogo server for sequence logo presentation ^57^.

Chemical drawings were performed using ChemDraw version 21.0.0 (PerkinElmer Informatics).

Structural figures were generated with Pymol version 2.4.2. (Schroedinger). Protein tunnels were visualized using the CAVER 3.0.3 algorithm ^58^ implemented in PyMol plugin. A probe radius of 0.8 Å was used to optimize its presentation. The approximate tunnel length of 27 Å was determined by the use of dummy atoms in 1 Å distances, as implemented in CAVER. Spatial residue differences from superimposed structures were determined by PDBeFold version 2.58 ^59^. Residue dihedral angles were calculated using MolProbity ^60^. Metal geometry was analyzed by CheckMyMetal ^24^. Changes in subunit arrangements were calculated using the PSICO module in PDBePISA, version 1.48 ^61^.

### Statistics and reproducibility

Relevant information on statistical data analysis is provided in relevant methods sections or legends of the respective figure panels. The numbers of experimental replicates are stated in the respective Table and Figure legends. Statistical analysis was performed using Origin 2020 (Origin Lab).

## AUTHOR INFORMATION

### Author Contributions

FF, RS and MW designed the project. FF, SSch performed the experiments. I.B. and G.B. performed the XRF experiment. FF, RS and MW analyzed the data. FF, RS and MW wrote the manuscript. RS and MW supported the project.

## ACKNOWLEDGMENT

We acknowledge technical support by the SPC facility at EMBL Hamburg and expert technical assistance by Jeannette Ueckert (University of Regensburg). The synchrotron data was collected at beamline P13, EMBL Hamburg at the PETRA III storage ring (DESY, Hamburg, Germany). Panagiotis Spatharas and David von Stetten are thanked for contributions on X-ray fluorescence measurements.

## ABBREVIATIONS

ADCS: aminodeoxychorismate synthase
AS: anthranilate synthase
Cho: chorismate
GAT: glutamine amidotransferase
GDH: glutamate dehydrogenase
Gln: glutamine
Gln-TE: glutamyl thioester
IMAC: immobilized metal affinity chromatography
ITC: isothermal titration calorimety
MST enzymes: menaquinone siderophore and tryptophan enzymes
PabA: ADCS glutaminase subunit
PabB: ADCS synthase subunit
SEC: size exclusion chromatography
XRF: X-ray fluorescence.

