## Supplement for "Activity regulation of a glutamine amidotransferase bienzyme complex by substrate-induced subunit interface expansion"

^$^Current address: European Molecular Biology Laboratory Grenoble, 38042 Grenoble Cedex 9, France

Corresponding authors:

**Supplement**

**Supplementary Table 1: X-ray crystallography data**

| **Protein** | **PabA/PabB** | **PabA/PabB** | **PabA/PabB** | **PabA/PabB** | **PabA** |
| --- | --- | --- | --- | --- | --- |
| **Chains** | A/D, B/C | A/D, B/C | A/D, B/C | A/D, B/C | A, B, C |
| **Added Ligands** |  | EDTA | Gln | Gln, Cho |  |
| **Found Ligands** | Zn^2+^, Trp | Trp | Gln-TE (A), Trp | Gln-TE (A), Cho, Trp |  |
| **PDB entry** | 8RP6 | 8RP2 | 8RP1 | 8RP0 | 8RP7 |
| **Data collection** | | | | | |
| Space group | P2_1_2_1_2_1_ | P2_1_2_1_2_1_ | P2_1_2_1_2_1_ | P2_1_2_1_2_1_ | P2_1_ |
| Cell dimensions  a, b, c (Å) | 78.5, 108.3, 179.8 | 78.9 109.4 178.9 | 79.4, 109.9, 174.3 | 80.1, 109.9, 175.6 | 36.8, 115.6, 63.5 |
| Resolution (Å) | 2.45 | 1.98 | 1.86 | 1.64 | 2.82 |
| R merge | 0.147 (1.699) | 0.103 (2.034) | 0.099 (1.090) | 0.120 (1.797) | 0.313 (1.486) |
| I/σ(I) | 11.9 (1.8) | 16.7 (1.67) | 16.7 (2.6) | 11.2 (1.2) | 6.8 (1.8) |
| CC 1/2 | 0.998 (0.806) | 0.999 (0.693) | 0.998 (0.853) | 0.997 (0.677) | 0.984 (0.513) |
| Completeness (%) | 99.88 (99.79) | 99.94 (99.95) | 99.96 (99.98) | 99.97 (99.99) | 98.54 (98.31) |
| Redundancy | 13.4 (13.5) | 13.2 (13.6) | 13.3 (13.5) | 13.2 (13.3) | 5.1 (5.1) |
| **Refinement** | | | | | |
| Resolution (Å) | 46.39-2.45 (2.54-2.45) | 44.9-1.98 (2.05-1.98) | 46.88-1.86 (1.93-1.86) | 46.6-1.64 (1.70-1.64) | 42.39-2.82 (2.92-2.82) |
| No. reflections | 57,801 | 108,389 | 128,391 | 189,813 | 12,411 |
| R work / R free | 0.204 / 0.254 | 0.200 / 0.239 | 0.167 / 0.201 | 0.169 / 0.199 | 0.211 / 0.268 |
| **No. atoms** | | | | | |
| Protein chains | 2 x PabA  2 x PabB | 2 x PabA  2 x PabB | 2 x PabA  2 x PabB | 2 x PabA  2 x PabB | 3 x PabA |
| Protein atoms | 10166 | 10183 | 10194 | 10215 | 4423 |
| Protein residues | 1285 | 1285 | 1287 | 1286 | 567 |
| Ligands | 25 | 69 | 159 | 103 |  |
| Water | 67 | 240 | 625 | 999 | 3 |
| **B factors** | | | | | |
| Protein | 63.2 | 50.9 | 36.7 | 35.8 | 56.1 |
| Ligand | 75.8 | 79.7 | 64.7 | 54.8 |  |
| Water | 48.1 | 45.1 | 41.5 | 43.5 | 36.3 |
| **R.m.s. deviations** | | | | | |
| Bond lengths (Å) | 0.007 | 0.009 | 0.011 | 0.011 | 0.009 |
| Bond angles (°) | 1.50 | 1.54 | 1.56 | 1.67 | 1.41 |
| **Ramachandran statistics (%)** | | | | | |
| Favored regions | 96.5 | 96.9 | 97.4 | 97.3 | 92.7 |
| Allowed regions | 3.4 | 2.9 | 2.6 | 2.7 | 6.1 |
| Outliers | 0.2 | 0.2 | 0.0 | 0.0 | 1.3 |

The highest resolution shell values are in parenthesis.

**Supplementary Table 2. Structural relations of representative ammonia-utilizing MST enzyme complexes**

| **Enzyme** | **Organism** | **PDB entry** | **Rmsd**  **[Å^2^]** | **N_res_ ^a^** | **N_PabA_ ^b^**  **[%]** | **Δϕ ^c^**  **[deg.]** | **Rmsd**  **[Å^2^]** | **N_res_ ^a^** | **N_PabB_ ^b^**  **[%]** | **Δϕ ^c^**  **[deg.]** |
| --- | --- | --- | --- | --- | --- | --- | --- | --- | --- | --- |
| **ADCS** | *Escherichia coli* | 8rp2 ^d^ | **PabA-based superposition** | | | | **PabB-based superposition** | | | |
|  | *Streptomyces venezuelae* | 8hx6 | 1.18 | 170 | 91 | 19.6 | 1.46 | 405 | 89 | 19.6 |
| **AS** | *Serratia marcescens* | 1i7s | 1.29 | 173 | 93 | 24.9 | 2.04 | 407 | 90 | 26.6 |
| **AS** | *Salmonella typhimurium* | 1i1q | 1.17 | 175 | 94 | 24.9 | 1.99 | 403 | 89 | 27.9 |
| **AS** | *Saccharolobus solfataricus* | 1qdl | 1.09 | 183 | 98 | 24.2 | 2.04 | 415 | 91 | 24.2 |
| **ADICS** | *Burkholderia lata* | 3r74 | 1.76 | 157 | 84 | 12.0 | 2.26 | 331 | 73 | 12 |

^a^ Number of matching residues with the target sequence.

^b^ Percentage of matching residues, using the PabA sequence (187 residues) or PabB sequence (455 residues) as reference.

^c^ Change in orientation of PabB/synthase target principal axes resulting from a PabA-based superposition and PabA/synthase target principal axis resulting from a PabB-based superposition, calculated with PSICO (for further details see Materials and Methods, **Supplementary Figure 7**). Comparison of each value pair revealing very similar data demonstrates the robustness of the calculation of the principal domain/subunit axes by PSICO.

^d^ ADCS coordinates of the glutaminase apo state (PDB ID: 8rp2) were used. All target structures were determined in the absence of glutaminase substrate, inhibitors or reaction ligands. The analysis demonstrates for representative structures of ammonia-utilizing MST enzyme complexes that despite close structural relationships at the level of separate glutaminase and synthase subunits/domains there is a substantial degree of variation in the orientation between the respective glutaminase and synthase domains/subunits, making an overall superposition of complete enzyme complexes unreliable or even impossible. Conformational changes at different catalytic states, as performed elsewhere in this contribution for different ADCS complexes, were not considered in this comparative analysis of members of the MST family.

**Supplementary Table 3: Gln-TE mediated PabA/PabB interface expansion and ADCS complex rearrangement**

| PDB entry | Gln-TE ^a^ | Cho/  Mg^2+^ ^a^ | Zn^2+, a^ | **Δϕ** [^o^] **^c^** | ASA  [Å^2^] ^d^ | N ^d^ | ΔG_calc_ [kcal/Mol]^d^ | K_D(calc)_ [μM] ^e^ | K_D(calc, apo) /_ K_D(calc, ligand)_ |
| --- | --- | --- | --- | --- | --- | --- | --- | --- | --- |
| 8RP6 |  |  | + |  | 904 | 10 | -7.7 | 2.2 |  |
| 8RP2 (EDTA) |  |  |  |  | 965 | 14 | -8.1 | 1.1 |  |
| 8RP1 (Gln ^b^) | + |  |  | 23.4 | 1312 | 28 | -10.6 | 0.017 | 129 |
| 8RP0 (Gln, Cho ^b^) | + | + |  | 23.2 | 1324 | 27 | -10.3 | 0.033 | 66 |

^a^ Ligand observed in the electron density of the crystal structure

^b^ Ligand added during crystallization

^c^ Change in orientation of PabB principal axis calculated with PSICO, resulting from PabA-based superposition and using ADCS coordinates with no reaction ligands added (8RP6) as reference (**Supplementary Figure 7**). Since the PabA/PabB arrangement in 8RP6 and 8RP2 (ADCS coordinates with EDTA added) is identical, the Δϕ angles are unchanged when 8RP2 is used as the reference.

^d^ ASA, accessible surface area; N, number of specific interactions between PabA and PabB subunits (hydrogen bonds, salt bridges), ΔG_calc_, solvation energy gain, as defined in PDBePISA, version 1.48.

^e^ K_D(calc)_ values were calculated by applying ΔG_calc_ = RT * lnK_D(calc)_

**Supplementary Table 4: Effects of PabA/PabB active site ligands on ADCS steady-state glutaminase activity ^a^**

| 0.2 mM Cho | 5 mM Mg^2+^ | 0.2 mM Zn^2+^ | 1 mM  EDTA | K_M_^Gln^  [mM] | k_cat_  [s^-1^] | k_cat_/K_M_^Gln^  [mM^-1^s^-1^] |
| --- | --- | --- | --- | --- | --- | --- |
| – | – | – | – | 0.09 ± 0.005 | 0.20 ± 0.003 | 2.35 ± 0.17 |
| – | + | – | – | 0.09 ± 0.01 | 0.22 ± 0.007 | 2.41 ± 0.47 |
| – | – | – | + | 0.08 ± 0.002 | 0.19 ± 0.001 | 2.42 ± 0.08 |
| – | + | – | + | 0.09 ± 0.005 | 0.23 ± 0.003 | 2.58 ± 0.17 |
| + | – | – | – | 0.07 ± 0.005 | 0.14 ± 0.003 | 1.98 ± 0.18 |
| + | + | – | – | 0.34 ± 0.03 | 0.37 ± 0.010 | 1.09 ± 0.13 |
| + | + | + | – | 0.31 ± 0.008 | 0.42 ± 0.003 | 1.33 ± 0.04 |
| + | – | – | + | 0.08 ± 0.002 | 0.15 ± 0.001 | 1.91 ± 0.07 |
| + | + | – | + | 0.27 ± 0.010 | 0.31 ± 0.003 | 1.13 ± 0.05 |
| + | + | + | + | 0.31 ± 0.0n8 | 0.38 ± 0.006 | 1.24 ± 0.09 |

To investigate the role of zinc ion binding in ADCS catalysis we additionally purified the ADCS complex under conditions in which 0.2 mM ZnCl_2_ was added to the expression medium to ensure an excess of zinc in the purified protein, since the addition of Zn^2+^ to the purified ADCS complex led to its precipitation. To measure ADCS glutaminase activity of these samples in the absence of Zn^2+^, samples were dialyzed against a buffer containing EDTA. We also measured ADCS glutaminase activity in the presence and absence of synthase substrate Cho and Mg^2+^**(Supplementary Figure 4a, Table 1)**. To this end, we could confirm previously established moderate changes on ADCS glutaminase activity by the presence of both Cho and Mg^2+^, leading to a three to fourfold increase in K_M_^Gln^ and an approximately twofold increase in k_cat_ of the ADCS glutaminase turnover **^1^**. These effects together resulted in a slight decrease in the ADCS glutaminase catalytic efficiency as measured by k_cat_/K_M_^Gln^. In contrast, we found no significant change in ADCS glutaminase activity, caused by the either enforced presence of Zn^2+^ through its addition to the expression medium or its removal by EDTA treatment on ADCS glutaminase activity. These results demonstrate that, despite the close proximity of the ADCS Zn^2+^ binding site to the glutaminase active site, the presence of Zn^2+^ does not play a role in ADCS glutaminase catalysis.

**Supplementary Table 5. PabA residues contributing to PabA/PabB interface, Zn^2+^ and Gln-TE binding**

| Residue | MC/SC ^a^ | IA ^b^ | Gln-TE ^c^ | Zn^2+, c^ | PabB, generic PabA/PabB interface | PabB, extended PabA/PabB interface |
| --- | --- | --- | --- | --- | --- | --- |
| aS10 | SC | HB |  |  |  | bK372 |
| aT12 | MC | HB |  |  | (bN312) | bN312 |
| aW13 | MC | HB |  |  | bN312 | bN312 |
| aN14 | SC | HB |  |  | bD308 | bD308 |
| aQ17 | SC | HB |  |  | bG315  bA318  (bG321) | bG315  bA318  bG321 |
| aY18 | SC | HB |  |  |  | bR311 |
| aR30* | SC | SB |  |  |  | bE119 |
| aG52 | MC | HB | X |  |  | bY210 |
| aC54* | MC | HB | X |  |  | bG207 |
| aC79 | SC | CB | X |  |  |  |
|  | SC |  |  | X |  |  |
| aL80 | MC | HB | X |  |  |  |
| aQ83 | SC | HB | X |  |  |  |
| aK103 | SC | SB |  |  | bD430 | bD430 |
| aY127* | SC | SB |  |  | bD308 | bL304  bD308 |
| aH128* | SC |  |  | X |  |  |
| aS129 | SC | HB |  |  |  | bL204 |
|  | MC | HB | X |  |  |  |
| aL130 | MC | HB | X |  |  |  |
| aH168 | SC |  |  | X |  |  |
| aE170 | MC | HB |  |  | bR311 | bR311 |
| aS171* | SC | HB |  |  | bD308 | bD308 |
| aI172 | MC | HB |  |  | bD308 | bD308 |
| Gln-TE | SC | HB |  |  |  |  |
|  | MC | HB |  |  |  | bG207  bY210 |

Asterisks: residues, which have been mutated for biophysical and biochemical characterization. Parentheses: Interactions not detected in all relevant structures.

^a^ MC, main chain; SC, side chain

^b^ IA, interaction; HB, hydrogen bond; SB, salt bridge. The interactions have been determined as defined in PDBePISA, version 1.48.

^c^ A distance cutoff of 3.5 Å was applied.

**Supplementary Table 6: Glutaminase activity regulation in MST GATs**

|  | **ADCS** | **AS** | **ADICS** |
| --- | --- | --- | --- |
| Glutaminase/Synthase interface expansion | x | x | X |
| Glutaminase/Synthase rearrangement | x |  |  |
| Glutaminase active site Zn^2+^ binding | x |  | X |
| Synthase Trp binding site | x | x |  |
| Trp feedback inhibition |  | x |  |

**Supplementary Table 7. PabB residues contributing to Cho, Mg^2+^ and Trp binding**

| Residue | MC/SC ^a^ | IA ^b^ | Cho ^c^ | Mg^2+, c, d^ | Trp ^c^ |
| --- | --- | --- | --- | --- | --- |
| bH35 | MC | HB |  |  | x |
| bS36 | SC | HB |  |  | x |
| bY43 | MC | HB |  |  | x |
| bR45 | MC | HB |  |  | x |
| bF46 | MC | HB |  |  | x |
| bP240 | MC | HB |  |  | x |
| bS242 | MC | HB |  |  | x |
| bG275 | MC | HB | x |  |  |
| bT276 | SC | HB | x |  |  |
|  | MC | HB |  | x |  |
| bK298 | SC | HB |  | x |  |
|  | MC | HB |  | x |  |
| bD299 | SC | HB |  | x |  |
| bE302 | SC | SB |  | x |  |
| bR410 | SC | HB | x |  |  |
| bG424 | MC | HB | x |  |  |
| bG426 | MC | HB | x |  |  |
| bE436 | SC | HB |  | x |  |
| bK443 | SC | HB | x |  |  |
| bE439 | SC | SB |  | x |  |

^a^ MC, main chain; SC, side chain

^b^ IA, interaction; HB, hydrogen bond; SB, salt bridge. Interactions were determined as defined in PDBePISA, version 1.48 (<https://www.ebi.ac.uk/pdbe/pisa/>).

^c^ A distance cutoff of 3.5 Å was applied.

^d^ The list also includes interactions with two structurally conserved water molecules that are coordinated by Mg^2+^ bound to the PabB synthase active site.

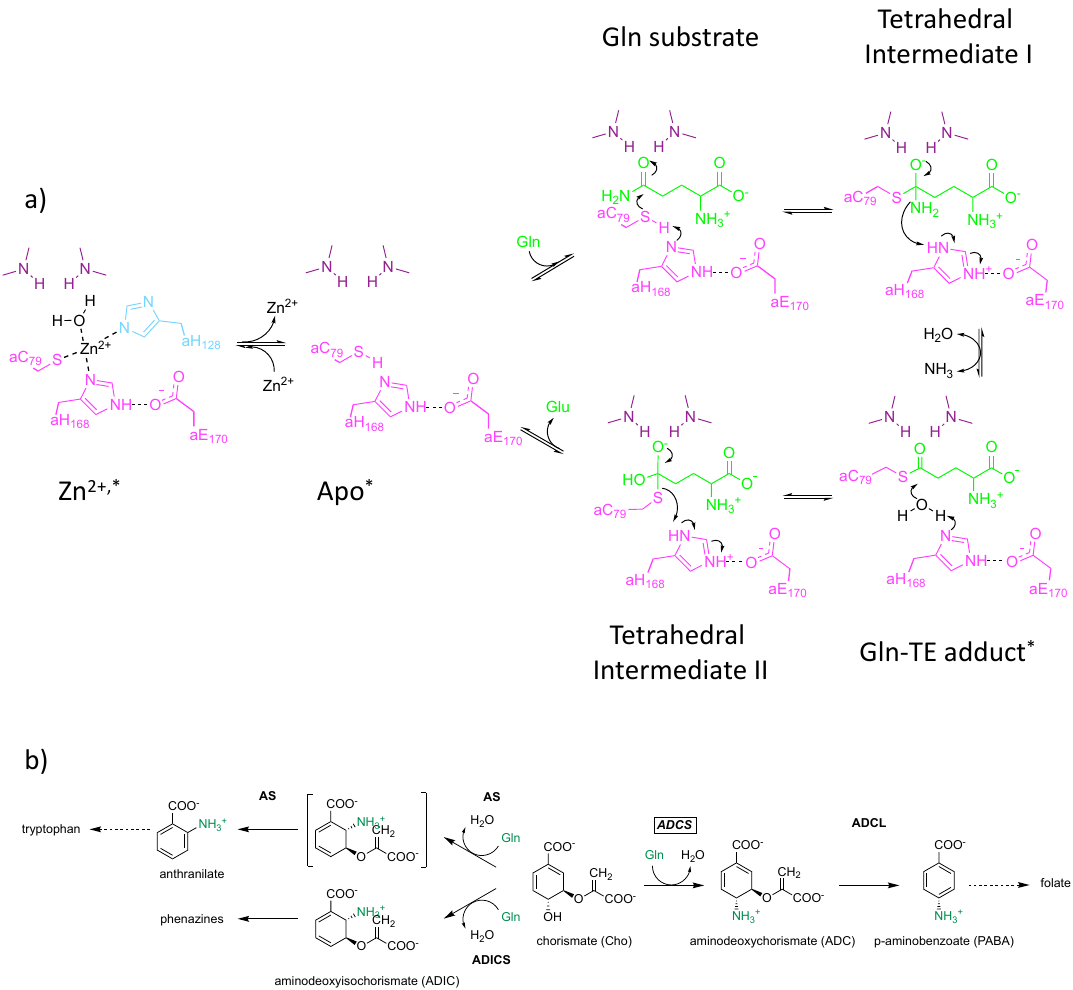

**Supplementary Figure 1: ADCS catalysis. a,** Glutaminase catalysis steps, adapted from **^2,3^**. Colors are as in **Figure 2**. ADCS glutaminase reaction states structurally investigated in this contribution are each marked with an asterisk. **b,** Synthase reactions catalyzed by related members of the MST family, which use glutaminase-produced ammonia as nucleophile and Cho as common synthase substrate. ACDS: aminodeoxychorismate synthase; AS: anthranilate synthase; ADICS: aminodeoxyisochorismate synthase; ACDL: aminodeoxychorismate lyase.

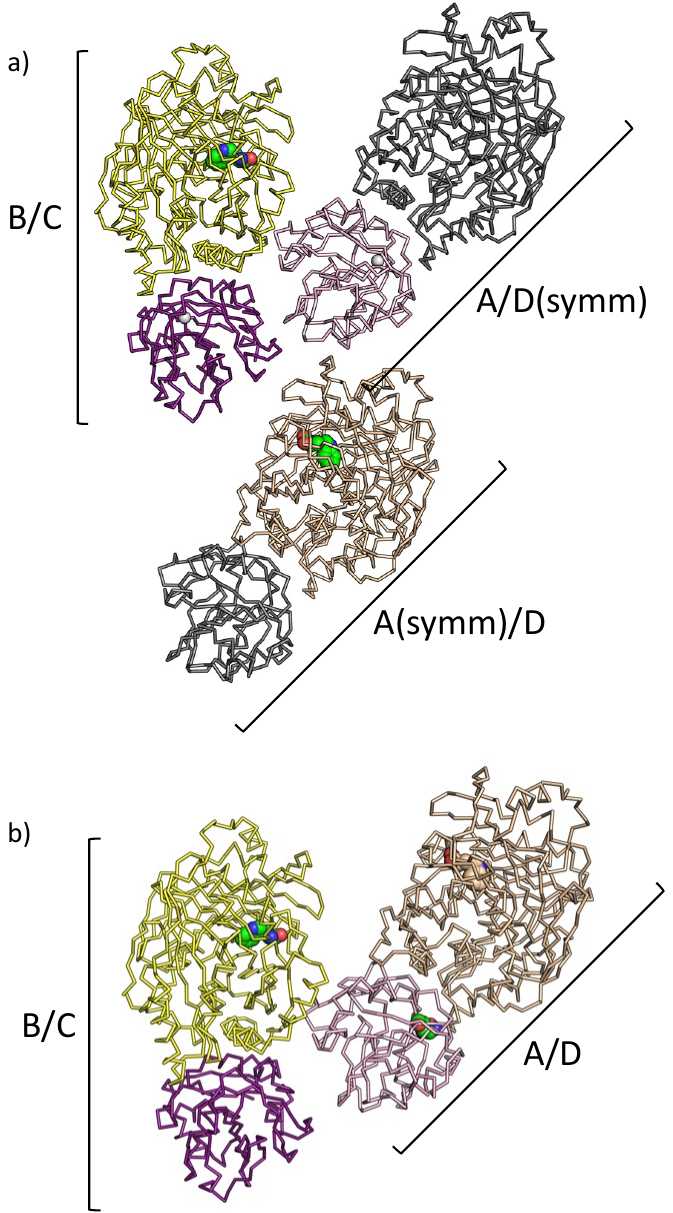

**Supplementary Figure 2. ADCS complexes in different crystal packing arrangements.** All crystal forms analyzed comprise two heterodimeric ADCS complexes **(*cf*. Supplementary Table 1).** **a,** in crystals used for ADCS structure determination without any reaction ligands or with added EDTA (PDB entries: 8RP6, 8RP2), the second ADCS complex is formed with symmetry-related subunits. Two of these complexes are illustrated, in which the symmetry-related subunit is shown in grey, while the other one is shown in colors related to the first ADCS complex, using the color conventions from **Figure 1**; **b,** in crystals used for ADCS structure determination with Gln added to the crystallization buffer (PDB entries: 8RP1, 8RP0), both biologically relevant ADCS complexes are situated within the same asymmetric unit. The main difference between the two ADCS crystal forms is in a change of the c axis of approximately 5 Å (***cf*. Supplementary Table 1**). The color codes are as in panel a. Bound Trp is marked by sphere presentation to facilitate orientation. For structure-based PabA/PabB interface analysis, we used only those PabA/PabB heterodimeric assemblies formed within the same asymmetric unit, to exclude any uncertainty in the measured distances due to experimental errors in the unit cell calculation **(Supplementary Table 3**).

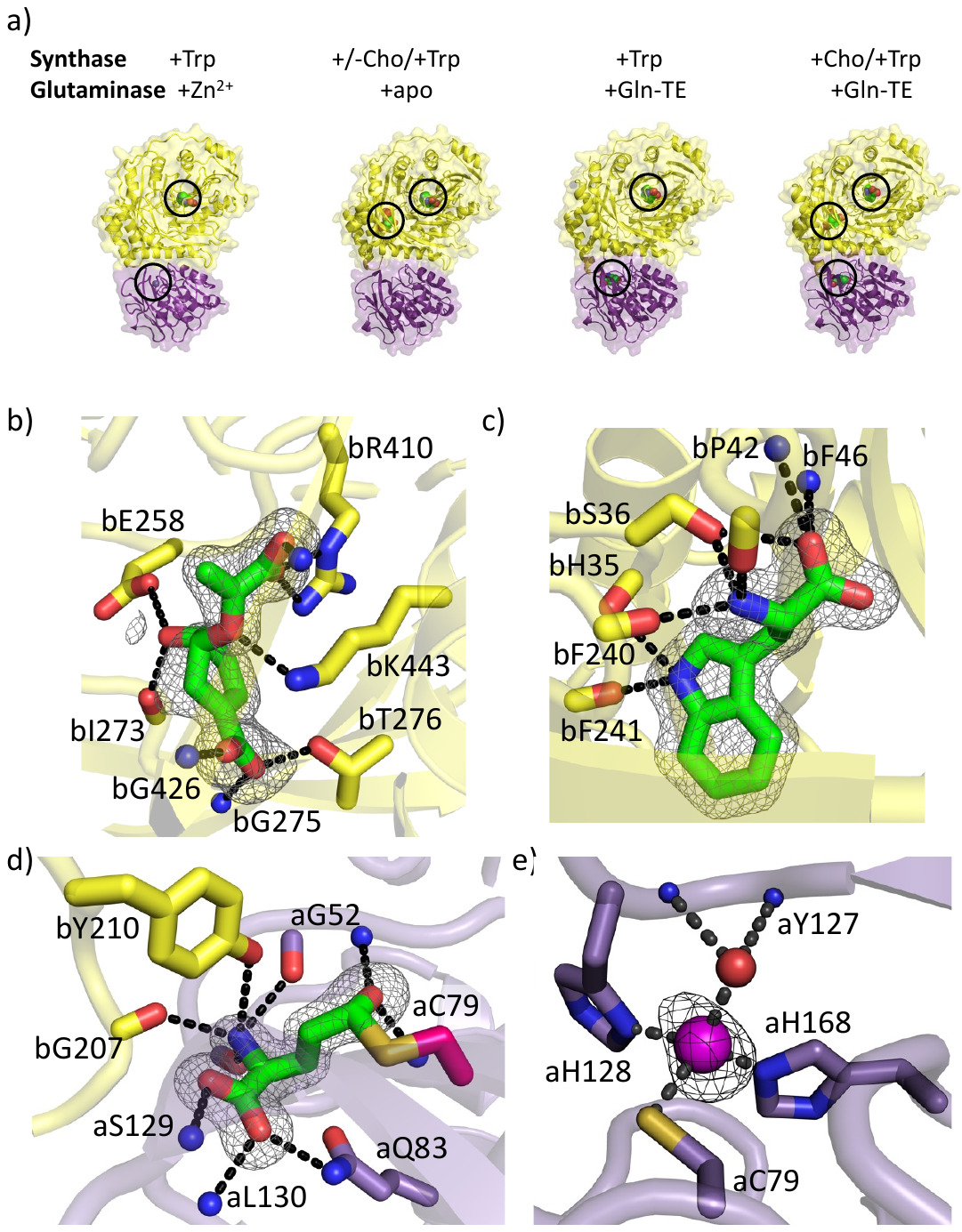

**Supplementary Figure 3:** **ADCS ligand binding sites, as evidenced by *2Fo-Fc* omit density at about 1 σ level. a,** Semi-transparent surface/cartoon representations of ADCS structures in different catalysis states, taken from **Figure 1b** as point of reference; **b**, Cho binding site (PDB entry: 8RP0, chain C) **(*cf*. Supplementary Figure 4, Supplementary Table 3);** **c**, Trp binding site (PDB entry: 8RP0, chain D) **(*cf*. Supplementary Table 3)**; d, Gln-TE binding site (PDB entry: 8RP0, chain A) **(*cf*. Figures 2c, 3a, Supplementary Table 3)**; e, Zn^2+^ binding site (PDB entry: 8RP6 chain B) **(*cf*. Figure 2a, Supplementary Table 3)**. Color codes are as in **Figure 2**. Selected interacting residues are shown in stick representation and are labeled. Hydrogen bonds are shown by dashed lines.

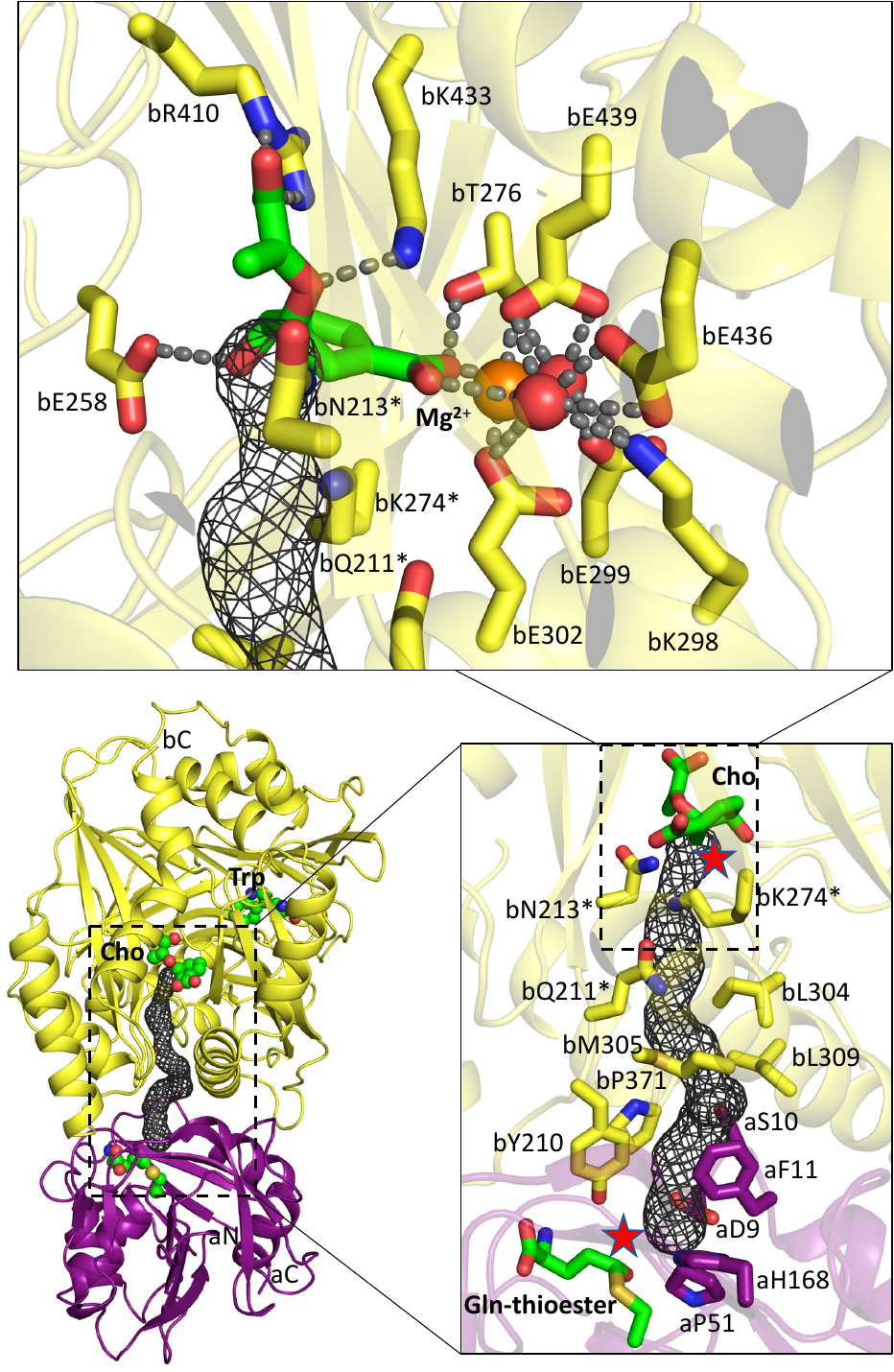

**Supplementary Figure 4: Structural details of ADCS glutaminase/synthase active sites-connecting ammonia tunnel.** Cartoon representation (lower left), showing ADCS complex formation and glutaminase/synthase active site connecting ammonia tunnel. Zoom on the ADCS ammonia tunnel (lower right). Residues interacting with dummy atoms of the ammonia tunnel are shown and labelled. The approximate position of ammonia leaving from Gln upon Gln-TE formation is indicated by a red asterisk. Zoom on the Cho/Mg^2+^ -binding site (upper panel), which also includes two structurally conserved water molecules (red spheres), tightly bound to Mg^2+^ (orange sphere) **(Supplementary Table 7)**. The exit of the glutaminase/synthase active sites connecting the ammonia tunnel is indicated by a grey mesh. Residues that are shown in both zoom panels are indicated by asterisks. To enhance insight into the underlying structures, the orientation of ammonia tunnel and the Cho/Mg^2+^ -binding site zoom is different, corresponding to a rotation around a vertical axis of about 45 degrees. PDB entry 8RP0, chains A and D, was used to generate the Figure.

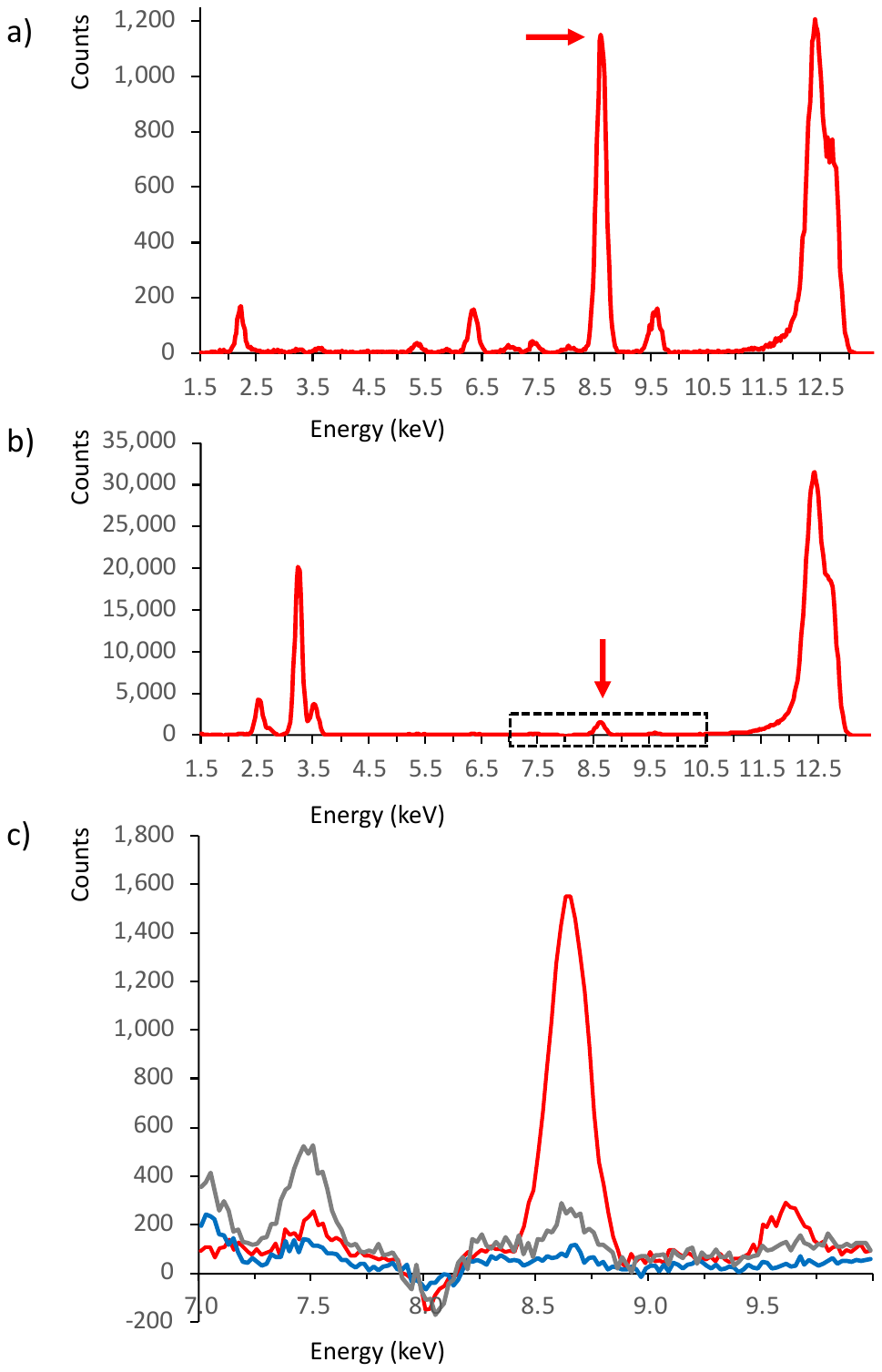

**Supplementary Figure 5. Evidence for Zn^2+^ in the glutaminase active site.** XRF scans of **a,** ADCS crystals without added reaction ligands; **b**, ADCS in solution without any reaction ligands added. Both spectra show a significant peak indicated by arrows at an energy that matches the K_α1_ edge of zinc at 8.64 keV. The large peak at 12.4 keV represents the energy of the primary X-ray beam. The other large peak of ADCS in solution is due to the K_α_ edge of K^+^ at 3.31 keV from the buffer used for purified ADCS; **c**, Zoom into the 7.0-10.5 keV energy segment (boxed in panel b) of ADCS complexes in solution: untreated ADCS, red (identical to panel b); EDTA-treated ADCS, blue; untreated ADCS aH128A variant, grey.

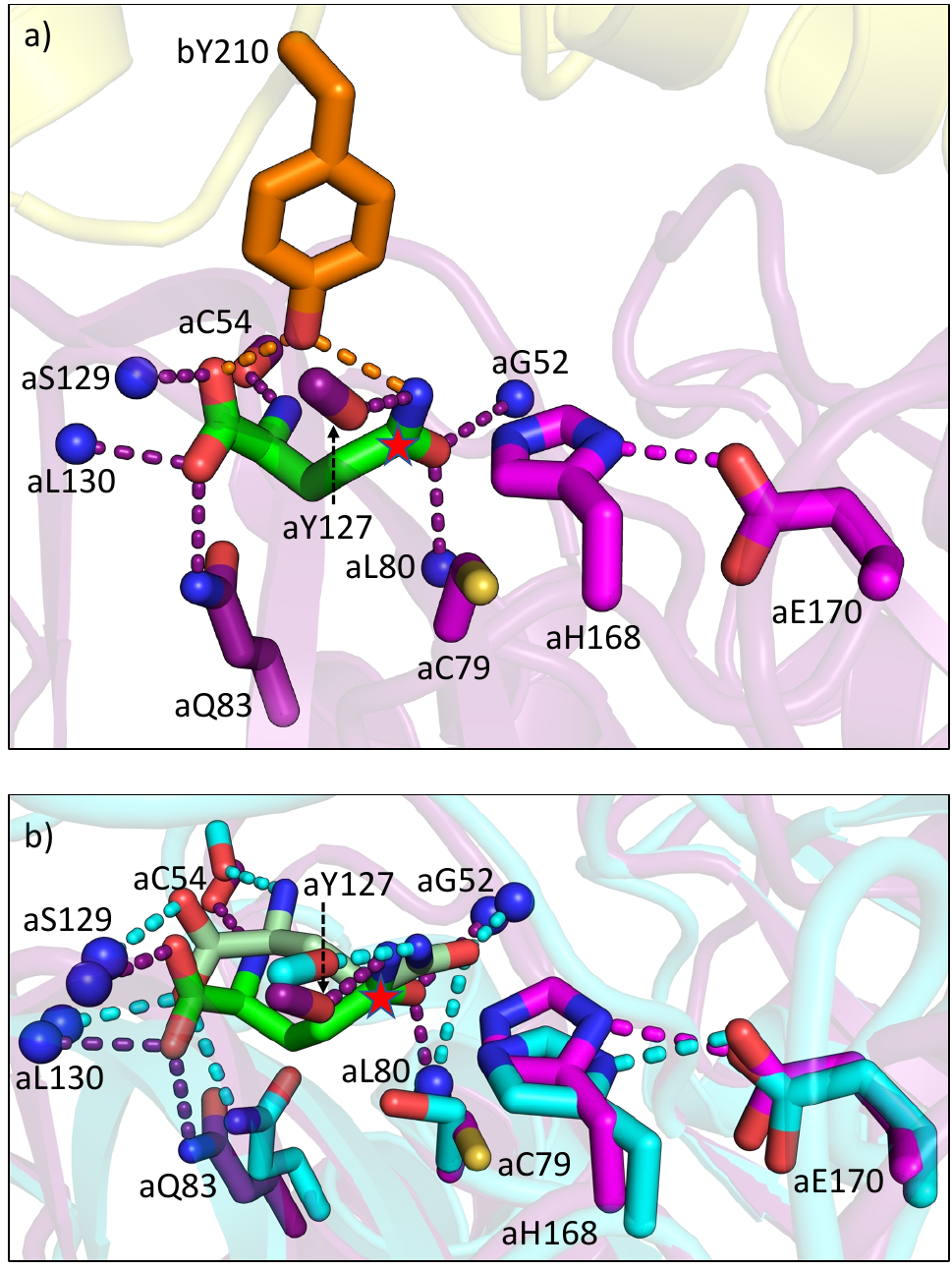

**Supplementary Figure 6. Modeling of Gln in the ADCS glutaminase active site.** **a,** Energy-minimized structural model of an ADCS Gln substrate complex (***cf*. Figure 2c**), based on the ADCS structure in the presence of Gln-TE **(Figure 3a, Supplementary Figure 3d)**; **b,** Superposition of the ADCS Gln substrate complex with the Gln-bound glutaminase domain of carbamoyl phosphate synthetase (PDB code: 1C3O; active site residues in cyan, Gln in pale green, r.m.s.d. = 1.13 Å). Structurally conserved hydrogen bond interactions with glutaminase active site residues are indicated by dashed lines in matching colors, demonstrating the close resemblance of the model of Gln binding in the two systems. There are further GAT glutaminase structures with active site-bound Gln (PDB codes: 2NV2, 2F2A, 3ZR4, 7AC8), which are structurally more distantly related. The Gln-TE carbon atom, where the substitution of the substrate amino group leading to TE formation and subsequent Gln-TE hydrolysis takes place, is marked with a red asterisk.

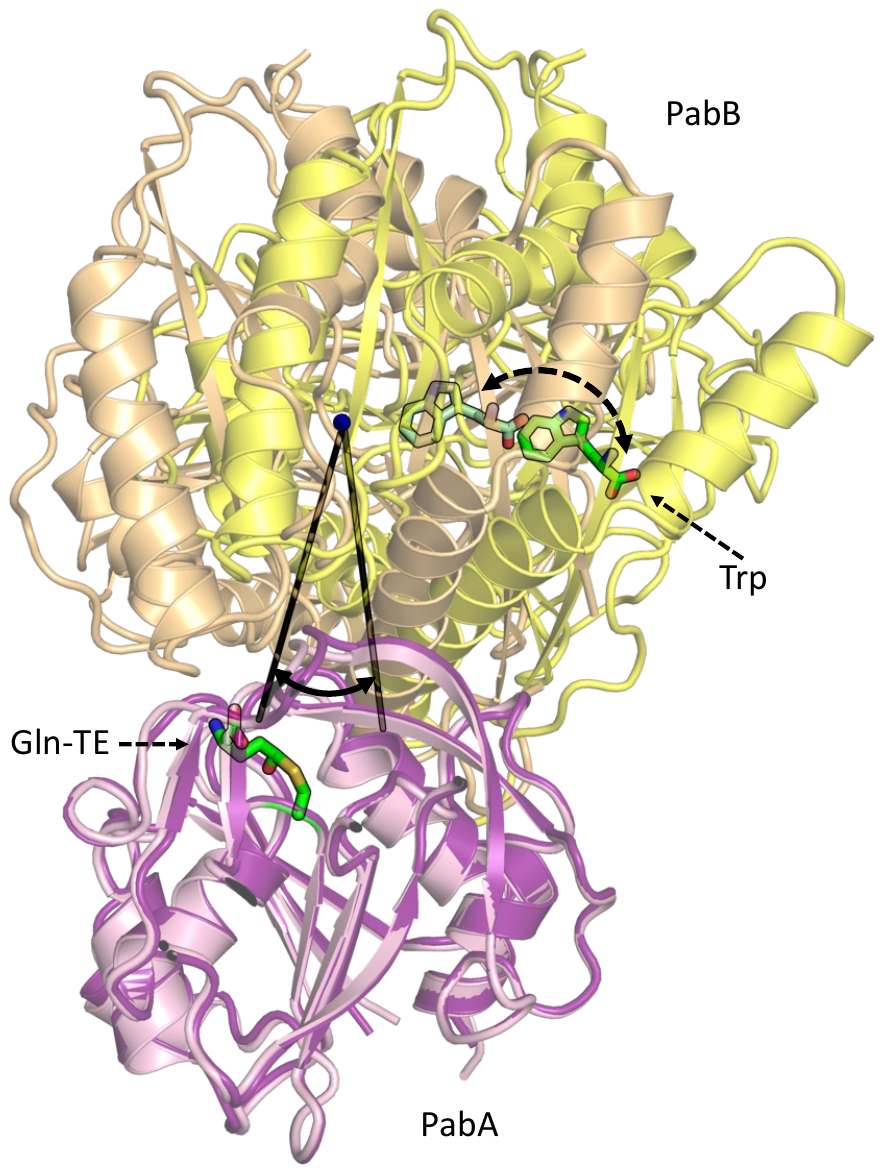

**Supplementary Figure 7. Analysis of the rigid-body rotation between the PabA and PabB subunits of the ADCS complex upon Gln binding.** PabA-based superposition of ADCS in the absence of reaction ligands (PDB entry: 8RP6 chains B and C) in colors as defined in **Figure 1** and Gln-TE bound ADCS (PDB entry: 8RP0 chains A and D) in related colors (PabA, light violet; PabB, light orange). The 23-degree rotation of the respective PabB subunits, as determined by PSICO/PyMol, is indicated. The Trp binding site in both structures is marked, to indicate the level of movement due to this rotational change. The bound Gln-TE in the glutaminase active site is also marked.

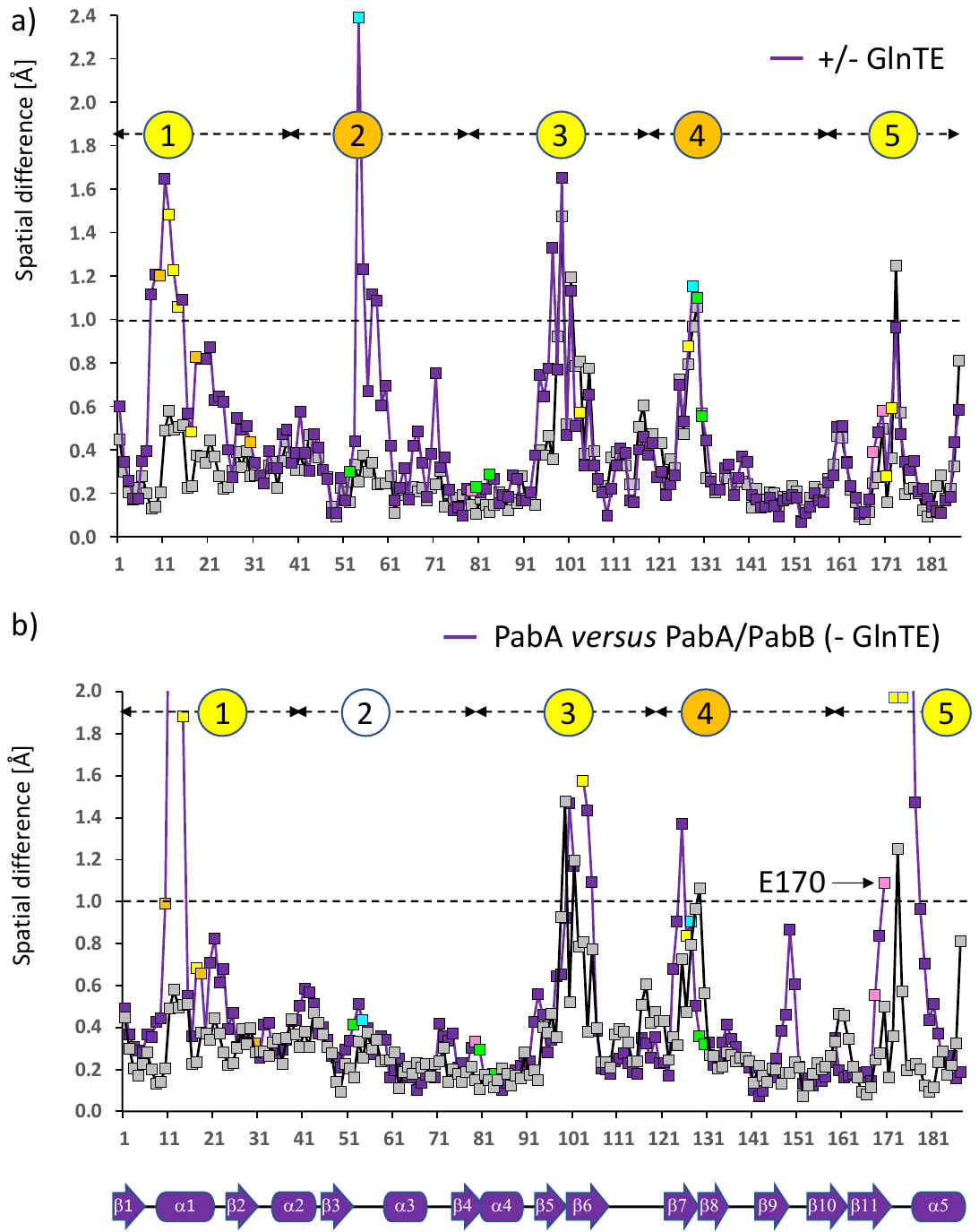

**Supplementary Figure 8. PabA conformational changes induced by the presence of Gln-TE and PabB:** **a,** R.m.s.d. plot indicating averaged spatial differences of PabA residues by superimposing structures of two ADCS heterodimeric complexes, where one PabA subunit is in the presence and the other one in the absence of Gln-TE (purple, PDB entries: 8RP1 chain A superimposed on 8RP6 chains A and B). **b,** R.m.s.d. plot indicating averaged spatial differences of PabA residues of the three separate PabA molecules in purple (PDB entry: 8RP7). For comparison, averaged spatial differences of PabA residues are also shown in grey, by superimposing structures of two ADCS heterodimeric complexes, in which both PabA subunits are devoid of Gln-TE (PDB entries: 8RP1 chain B superimposed on 8RP6 chains A and B). Residues that are involved in different parts of the PabA/PabB interface are colored as in the previous figures**.** The plots indicate five highly flexible sequence segments with spatial differences exceeding 1 Å, indicated by a dashed line. These segments are numbered 1-5 and shown in colors, representing their roles in different parts of the PabA/PabB interface (*cf*. **Figure 4**). The sequence positions of the secondary structural elements found in the structures of PabA are indicated below in linear cartoon representation. In contrast, we did not detect any degree of conformational changes in any of the PabB segments involved in the PabA/PabB interface **(Supplementary Figure 12)**, demonstrating that the glutaminase active site rigidification due to ADCS assembly is restricted to PabA.

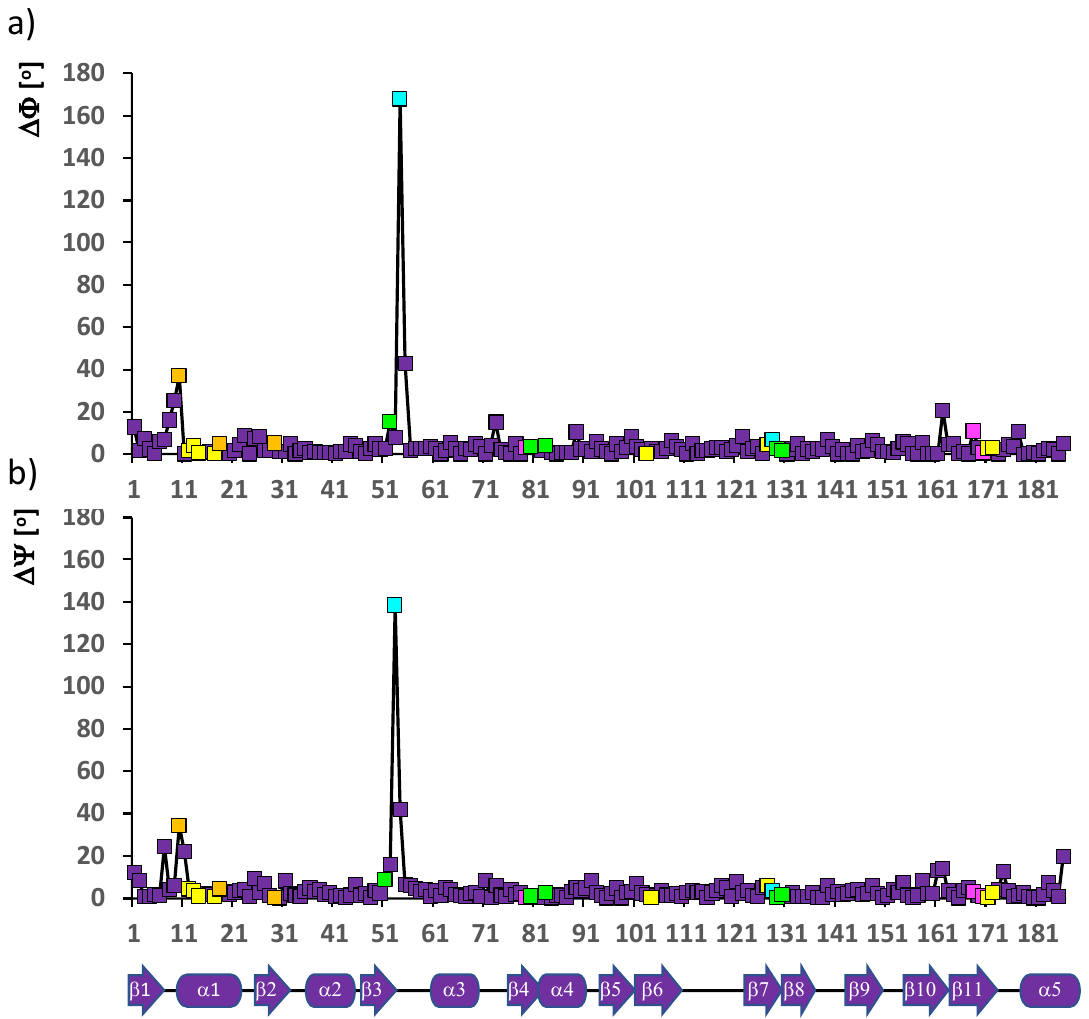

**Supplementary Figure 9. PabA dihedral angle changes induced by the presence of Gln-TE. a,** Absolute values of dihedral Φ angle differences of PabA subunits in the presence and absence of Gln-TE; **b,** absolute values of dihedral Ψ angle differences of PabA subunits in the presence and absence of Gln-TE.

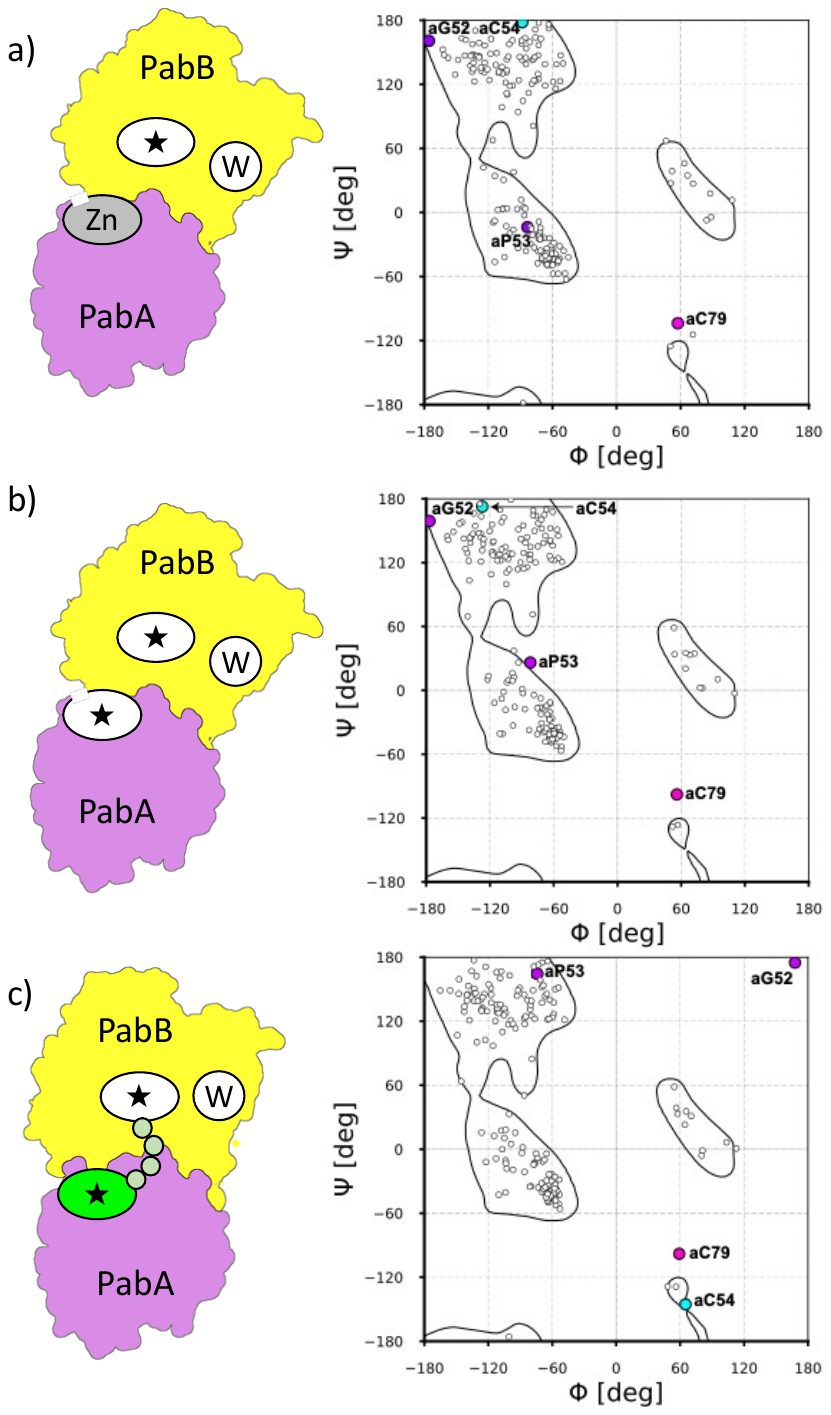

**Supplementary Figure 10. Ramachandran plot of PabA. a,** Plot in presence of Zn^2+^ (PDB code: 8RP6 chain B), **b,** in absence of any glutaminase active site ligands (PDB code: 8RP2 chain B) or **c**, with bound Gln-TE (PDB code: 8RP0 chain A). Residues aG52, aP53, aC54 and aC79 are highlighted. Color codes are as in **Figure 2ff.** Corresponding schematic ADCS PabA/PabB arrangements are shown on the left (***cf*. Figure 1**).

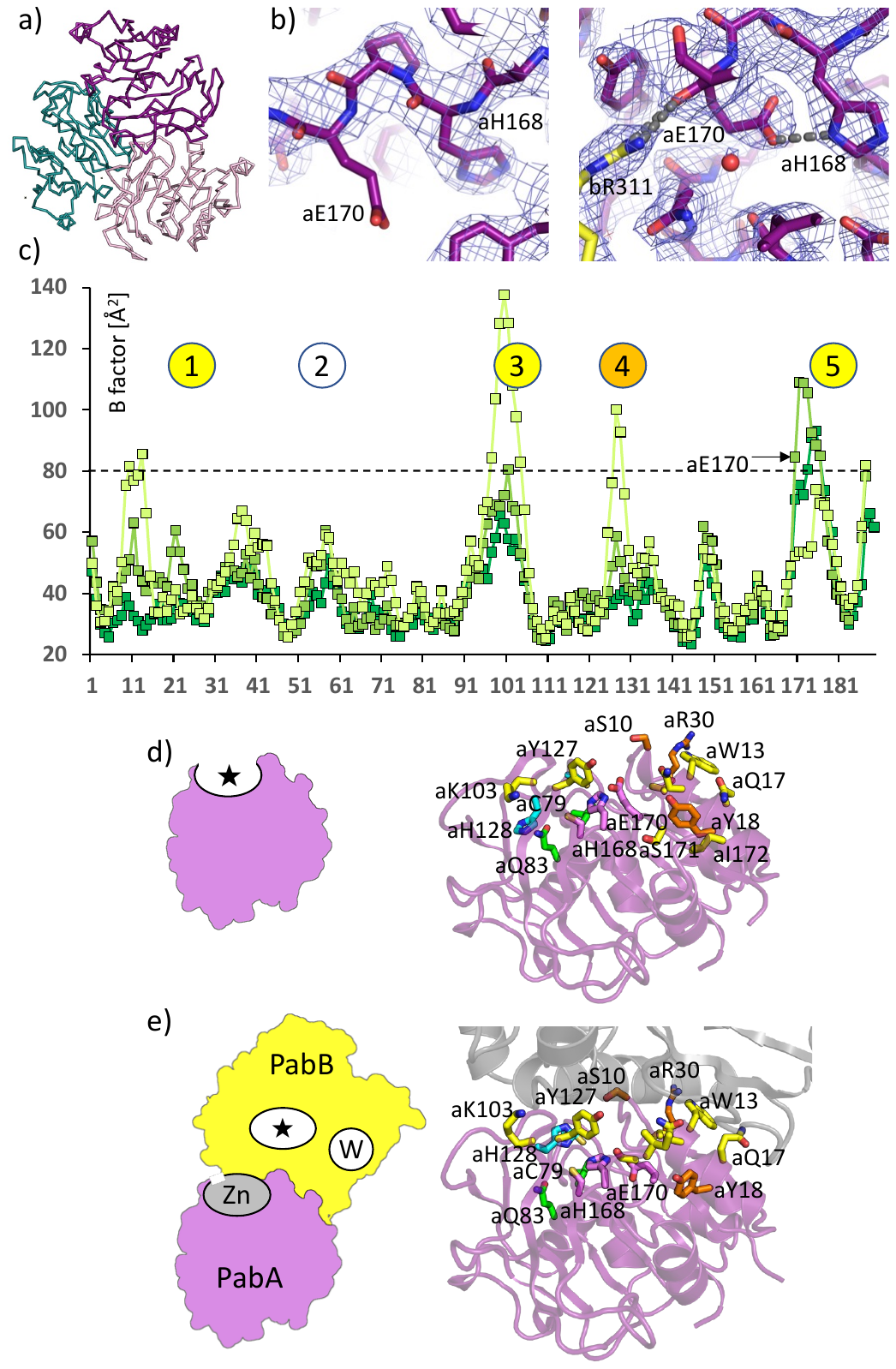

**Supplementary Figure 11. Increased conformational flexibility of the glutaminase active site in separate PabA subunits. a,** Arrangement of three separate PabA subunits within the crystal lattice (PDB entry: 8RP7). **b**, Left panel: Loose arrangement of catalytic triad residues aH168 and aE170 in separate PabA (PDB entry: 8RP7, chain A). Right panel: Catalytically competent arrangement of catalytic triad residues aH168 and aE170 in ADCS complexes (PDB entry: 8RP2, chains B and C), for comparison **(*cf*. Figures 2d, 3a)**. Hydrogen bonds connecting the side chains of aH168 and aE170, and the main chain carbonyl group of aE170 with the side chain of bE311 from the PabB subunit, are shown by dashed lines in grey. The structural models are displayed within the corresponding *2Fo-Fc* density maps used for modeling, to highlight the observed rigidification of the catalytic triad residues by the presence of PabB. In separate PabA, the high flexibility of the glutaminase active site leads to variable catalytic triad (aC79, aH68, aE170) arrangements, in which especially aE170 does not consistently adopt a position that allows to play its functional role as a relay of aH168 to enhance its function as nucleophile during glutaminase catalysis **(Figures 2c-d, 3a).** In contrast, in the structures of ADCS complexes there is a specific interaction between the main chain carbonyl group of aE170 and bR311, indicating an additional crucial role of PabB to generate a catalytically competent conformation of the glutaminase catalytic triad, irrespective of the presence of Gln **(Figure 4). c,** Residual mobility plots derived from refined crystal structures of the three separate PabA molecules (PDB entry: 8RP7) in different green shades. High mobility areas are numbered and match the structurally most divergent regions in PabA **(Supplementary Figure 8).** Cartoon representation of **d,** the separate PabA subunit structure (PDB entry: 8RP7, chain A) and **e,** the PabA subunit of the ADCS complex in complex with PabB, in the absence of Gln-TE (PDB entry: 8RP6). Color codes: PabA, violet; PabB, grey. PabA residues involved in the PabA/PabB interface in ADCS complex structures are shown in complementary PabB residue colors **(*cf*. Figure 4c, d).** Corresponding schematic PabA and PabA/PabB ADCS complex arrangements **(*cf*. Figure 1c)** are shown on the left.

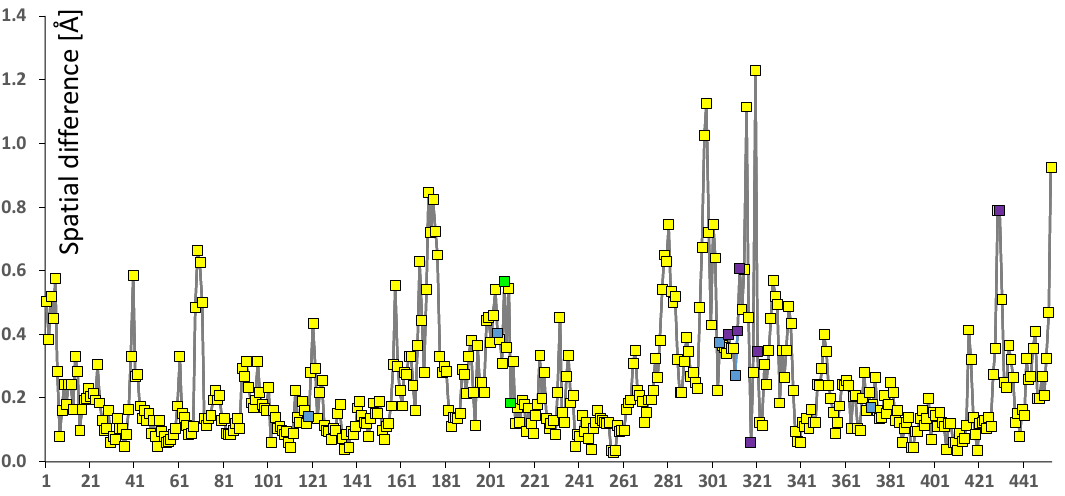

**Supplementary Figure 12**. **The PabB mobility is not significantly affected by Gln-TE binding to the ADCS complex.** R.m.s.d. plot indicating averaged spatial differences of PabB residues by superimposing structures of two heterodimeric ADCS complexes, where one interacting PabA subunit is in the presence and the other one in the absence of Gln-TE (PDB entries: 8RP1 chain D superimposed on 8RP6 chains C and D). A similar plot for the PabA sequence is shown in **Supplementary Figure 8a.** Note that most PabA/PabB interface residues from the PabB sequence do not reveal large scale spatial deviations, contrary to observations for PabA**.** Colors deviating from yellow are used for PabB residues contributing to the generic PabA/PabB interface in dark violet, contributing to the Gln-TE induced extended PabA/PabB interface in blue, and directly interacting with Gln-TE (when present) in green.

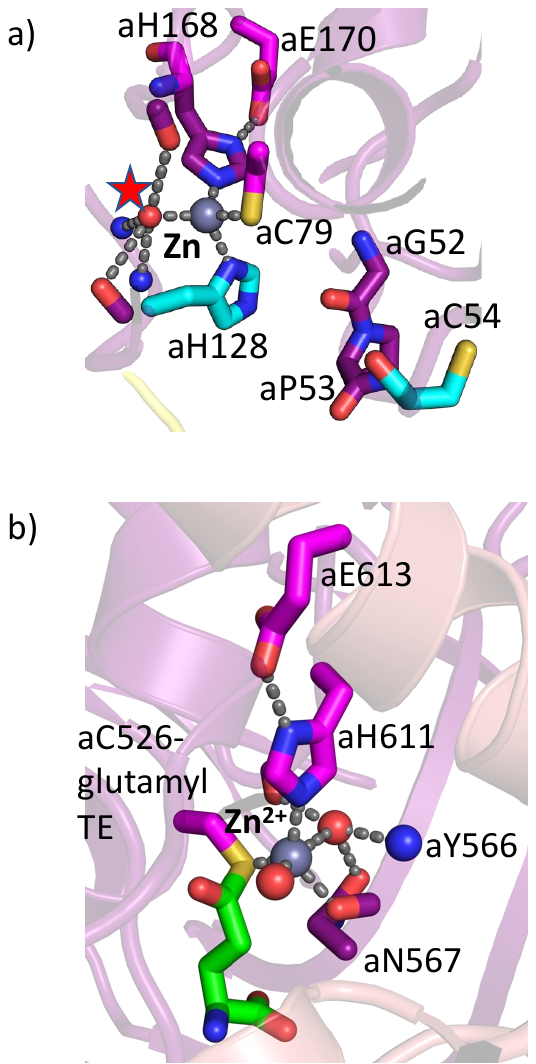

**Supplementary Figure 13**. **Structurally conserved Zn^2+^ in the glutaminase active sites** of **a**) ADCS **(*cf*. Figure 2a)** and **b**) ADICS (PDB code: 3R75). Contrary to ADICS, in which Zn^2+^ binding was observed in the presence of Gln-TE and could potentially act as a Lewis acid to enhance catalysis **^4^**, in ADCS we found Zn^2+^ only in the absence of Gln-TE, coupled with a swinging motion of aH128 and essentially blocking the ADCS glutaminase active site for substrate access **(Figure 2).** In ADICS, no comparable conformational change of any residue next to the glutaminase active site upon Zn^2+^ binding was observed. Interestingly, in both GATs Zn^2+^ appears to be extracted from the expression host, as no zinc ions were added during the purification and crystallization procedure ^5^. In both systems, zinc disappears from the glutaminase active site upon EDTA treatment, which is associated with no or only minor changes in glutaminase turnover **(Supplementary Table 4) ^5^.**

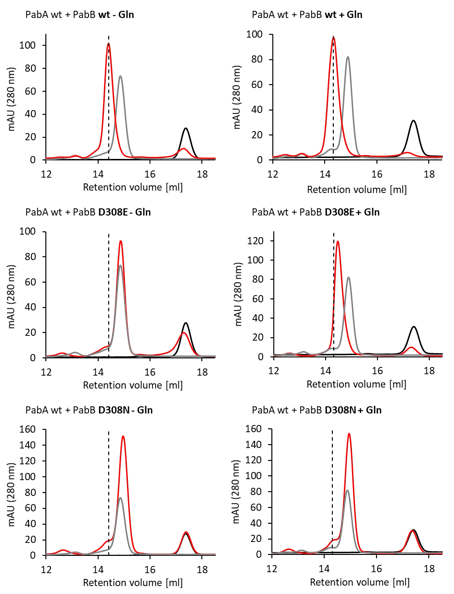

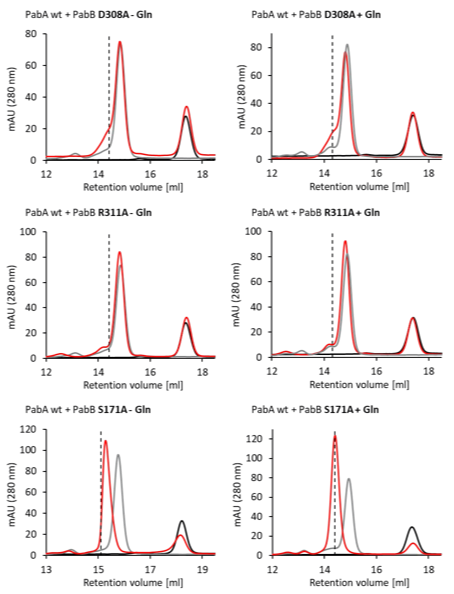

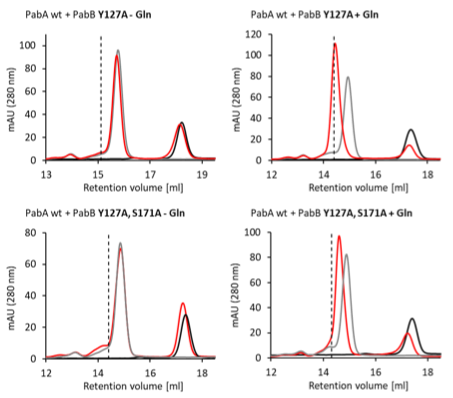

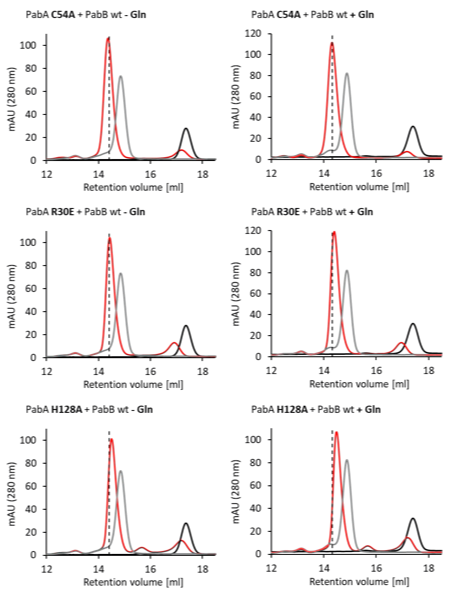

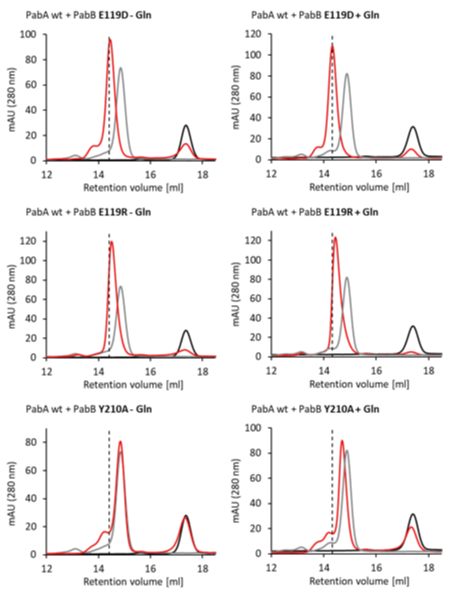

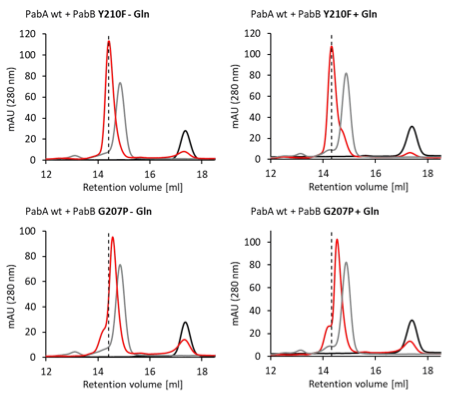

**Supplementary Figure 14.** **Elution profiles of analytical SEC to analyze Gln-dependent PabA/PabB complex formation.** Analytical SEC was performed on a Superdex 200 column equilibrated with 50 mM Tris/HCl pH 7.5, 50 mM KCl, 5 mM MgCl_2_, 2 mM DTT and, where indicated, 5 mM Gln. Dashed line, retention volume of the *wt* PabA/PabB complex; black, *wt* PabA; grey, *wt* PabB; red, ADCS variant as indicated above each graph.
